# ABCA7-dependent Neuropeptide-Y signalling is a resilience mechanism required for synaptic integrity in Alzheimer’s disease

**DOI:** 10.1101/2024.01.02.573893

**Authors:** Hüseyin Tayran, Elanur Yilmaz, Prabesh Bhattarai, Yuhao Min, Xue Wang, Yiyi Ma, Nastasia Nelson, Nada Kassara, Mehmet Ilyas Cosacak, Ruya Merve Dogru, Dolly Reyes-Dumeyer, Joseph S Reddy, Min Qiao, Delaney Flaherty, Andrew F. Teich, Tamil Iniyan Gunasekaran, Zikun Yang, Giuseppe Tosto, Badri N Vardarajan, Özkan İş, Nilüfer Ertekin-Taner, Richard Mayeux, Caghan Kizil

## Abstract

Alzheimer’s disease (AD) remains a complex challenge characterized by cognitive decline and memory loss. Genetic variations have emerged as crucial players in the etiology of AD, enabling hope for a better understanding of the disease mechanisms; yet the specific mechanism of action for those genetic variants remain uncertain. Animal models with reminiscent disease pathology could uncover previously uncharacterized roles of these genes. Using CRISPR/Cas9 gene editing, we generated a knockout model for *abca7,* orthologous to human *ABCA7 –* an established AD-risk gene. The *abca7*^+/-^ zebrafish showed reduced astroglial proliferation, synaptic density, and microglial abundance in response to amyloid beta 42 (Aβ42). Single-cell transcriptomics revealed *abca7*-dependent neuronal and glial cellular crosstalk through neuropeptide Y (NPY) signaling. The *abca7* knockout reduced the expression of *npy, bdnf* and *ngfra*, which are required for synaptic integrity and astroglial proliferation. With clinical data in humans, we showed reduced *NPY* in AD correlates with elevated Braak stage, predicted regulatory interaction between *NPY* and *BDNF*, identified genetic variants in *NPY* associated with AD, found segregation of variants in *ABCA7, BDNF* and *NGFR* in AD families, and discovered epigenetic changes in the promoter regions of *NPY, NGFR* and *BDNF* in humans with specific single nucleotide polymorphisms in *ABCA7*. These results suggest that ABCA7-dependent NPY signaling is required for synaptic integrity, the impairment of which generates a risk factor for AD through compromised brain resilience.

Graphical abstract

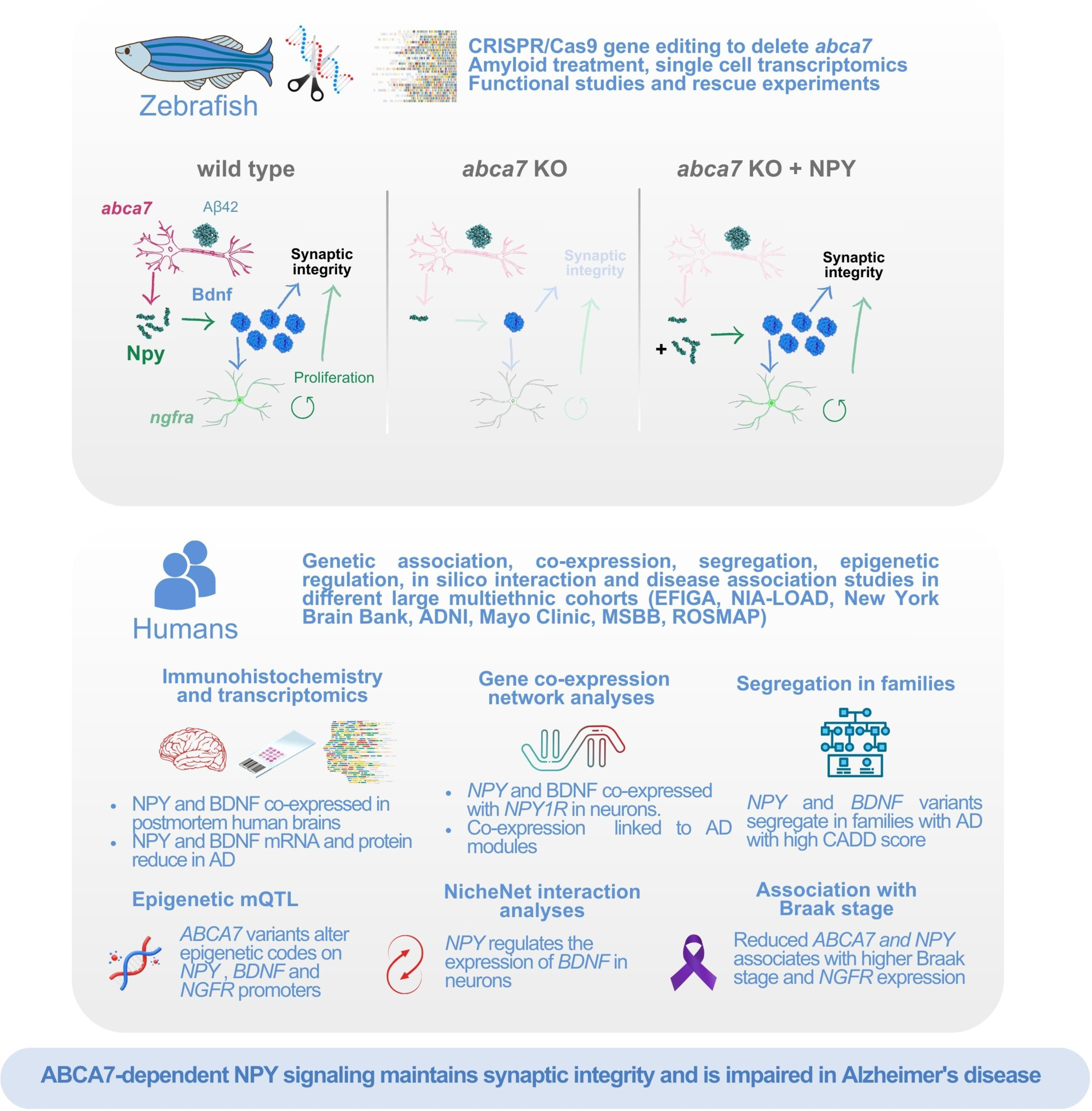

## Introduction

Alzheimer’s disease (AD), a complex and progressive neurodegenerative disorder, continues to present a formidable challenge to the society. This currently incurable condition, characterized by cognitive decline, particularly memory loss, and impaired daily functioning, has spurred large research efforts aimed at elucidating its underlying mechanisms. Among the multifaceted factors implicated in AD etiology, genetic variations have emerged as key players, offering insights into potential avenues of exploration ^1-5^. Despite the growing body of genetic evidence, the specific functions and mechanisms through which these genetic variations exert their influence remain a subject of ongoing investigation ^1,2, 5^. While genome wide association studies have provided significant data linking certain candidate genes to AD susceptibility, deciphering the mechanistic roles of these genes demands scrutiny into their functional significance, which would be facilitated by disease-mimetic representative animal models. Organisms that share conserved genetic elements with humans, offer a valuable platform for robust functional analysis. We developed a zebrafish model of amyloid toxicity, which shows strong parallels in the cellular and molecular mechanisms of the disease, including vascular perturbations, neurodegeneration, and immune reaction ^6-13^. This model has enabled functional investigation of AD genetic variants ^14-16^ and development of new active and potent pharmacological agents ^6,17, 18^. One of the key strengths of zebrafish is its remarkable neuro-regenerative ability after amyloid toxicity, which involves activation of molecular mechanisms that govern astroglial proliferation ^19-24^. Translational approaches that activate the molecular mechanisms in zebrafish to counteract amyloid toxicity through enhanced neurogenesis led to reduction of AD pathology in this model system ^25-29^. Therefore, zebrafish can serve as a useful *in vivo* translational and functional research tool for investigating the synaptic integrity and resilience mechanisms in AD.

Multiple studies have indicated that both common and rare variants in *ABCA7* are strongly and consistently associated with AD risk and endophenotypes across different ethnic groups ^30-39^. *ABCA7* encodes a transmembrane transporter with a demonstrated impact on lipid transport, a process integral to cellular membrane dynamics and overall homeostasis ^30,33,40-43^. By facilitating the movement of lipids across cellular membranes, ABCA7 influences a range of cellular processes, including those vital for neuronal function and synaptic plasticity. ABCA7 also modulates Aβ metabolism, a process central to the formation of the characteristic neuritic plaques observed in AD ^44,45^. These insights collectively underscore the potential significance of ABCA7 in orchestrating cellular mechanisms involved in AD pathogenesis. However, ABCA7 may have other roles and cellular mechanistic involvement in AD pathology that are yet uncharacterized, which we sought to uncover in this study that combined experiments and data from zebrafish and humans.

## Results

### *abca7* knockout affects the astroglial proliferation, synaptic density and microglia following amyloid toxicity

To determine the effects of heterozygous deletion of *abca7* in the zebrafish brain, we generated a truncated version of the gene by deleting 44.5 kilobases of genomic region between the Exon 13 and Exon 46 with CRISPR/Cas9-based gene editing (Figure 1A, B; Supplementary Table 1). This genomic deletion removes the protein domains after the second transmembrane domain including the ATPase and transporter domains, generating a null allele, similar to the human frameshift variation that is associated with AD risk ^33,46-48^. This deletion is genetically stable in later generations (Figure 1C) and reduces the expression of *abca7* gene (Figure 1D). We found that *abca7*^+/-^ animals are viable, fertile and do not show any phenotypes as larvae and adults. To investigate the adult brains of *abca7*^+/-^ zebrafish, we performed immunohistochemical stains for astroglial (GS), microglial (L-Plastin), synaptic (SV2), and proliferation (PCNA) markers in the presence or absence of Aβ42 toxicity (Figure 1E, F). Compared to wild type animals, *abca7*^+/-^ knockouts did not change astroglial proliferation, microglial numbers, or synaptic density. However, after Aβ42 was introduced through cerebroventricular microinjection, *abca7*^+/-^ knockouts displayed a significant reduction in astroglial proliferation, reduced the synaptic density, and increased the number of microglia (Figure 1G, Supplementary Data 1). These results suggest that *abca7* gene function is related to Aβ42-induced astroglial proliferation, synaptic integrity, and immune system reaction in zebrafish brain.

**Figure 1:**
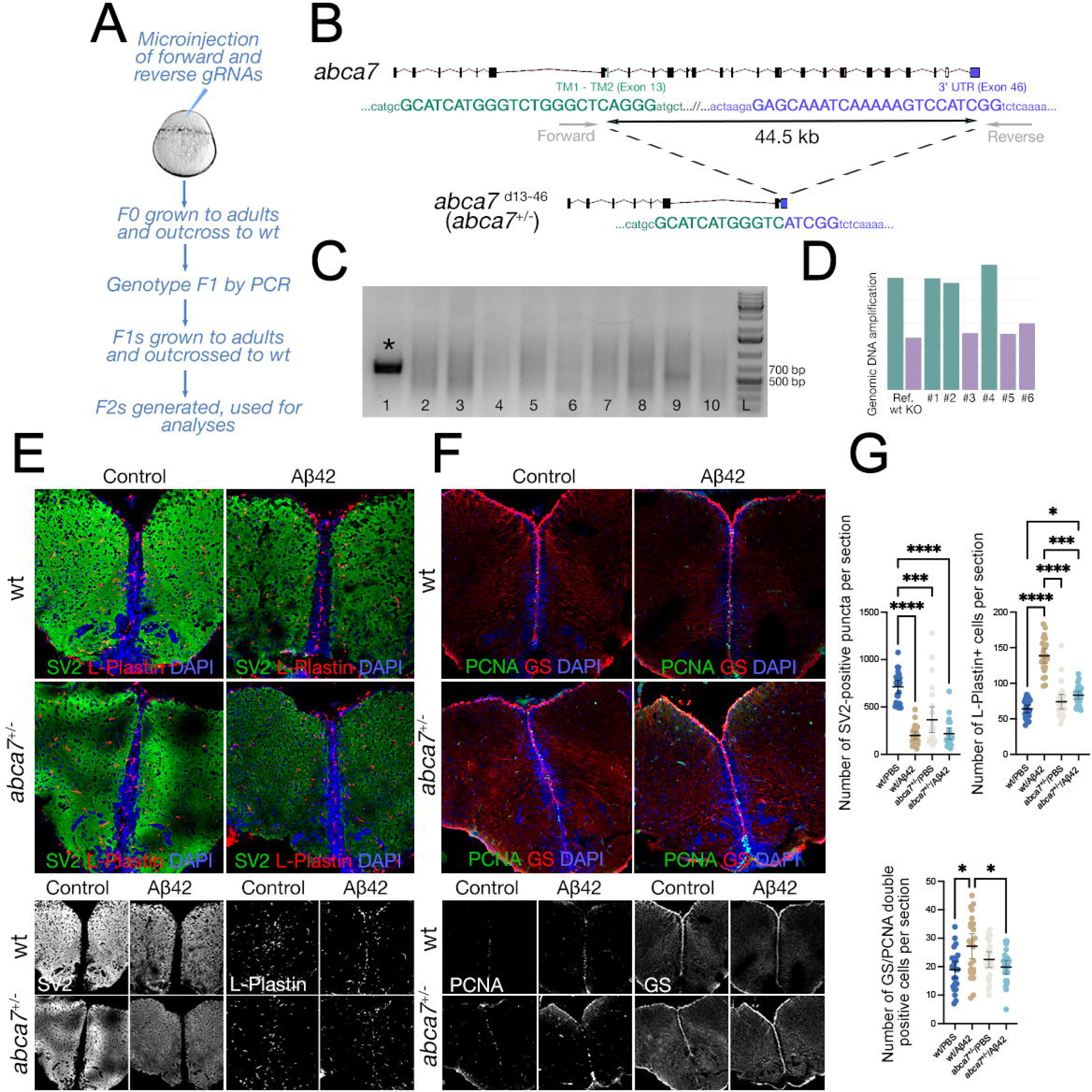
***abca7* is required for synaptic integrity, microglial prevalence and astroglial proliferation in zebrafish.** (A) Schematic view of generating *abca7* gene editing. (B) 44.5 kilobase deletion in *abca7* gene. (C) Representative genotyping PCR gel where a positive band indicates the deletion. (D) Genotyping results with genomic DNA quantitative PCR. Heterozygous deletions show reduced amplification. (E) Immunofluorescence (IF) for SV2 (green) and L-Plastin (red) with DAPI counterstain in wild type and *abca7*^+/-^ with and without Aβ42. Black and white panels indicate individual fluorescent channels. (F) IF for PCNA (green) and GS (red) with DAPI counterstain in wild type and *abca7*^+/-^ with and without Aβ42. Black and white panels indicate individual fluorescent channels. (G) Quantification of SV2-positive synaptic puncta, number of microglial cells, and number of proliferating astroglia.

### *abca7* is required for neuropeptide Y (*npy*) expression as a response to amyloid toxicity

We performed single cell transcriptomics in wild type/PBS-injected, wild type/Aβ42-injected, *abca7*^+/-^/PBS-injected, and *abca7*^+/-^/Aβ42-injected zebrafish (Figure 2A) to investigate the molecular basis of the *abca7*-dependent alterations. After quality control for mitochondrial gene ratios, abundance of ribosomal gene expression, distribution of the number of genes per cell, and number of reads per cell (Figure 2B, C), we determined the main cell type markers to categorize the 36 cell clusters (Figure 2D), which include 11 neuronal, 4 astroglial, 4 microglial, 3 immature neuronal, 2 immune, 2 vascular smooth muscle cell, 2 endothelial, 2 oligodendrocyte progenitor, 3 uncategorized, 1 neuroblast, 1 oligodendrocytic, and 1 pericytic cell cluster (Figure 2D, Supplementary Figure 1). In total, 50,691 cells passed the quality criteria and were clustered in the final UMAP (Figure 2E). 57.82% of the cells were neurons, followed by 10.35% astroglia, and 5.88% microglia (Figure 2E). The number of cells sequenced in all clusters (Figure 2F) and the number of cells sequenced per cell type in four samples combined and individually (Figure 2G) passed the power analyses by SCOPIT ^49^, therefore we proceeded with the analyses of cell clusters without exclusions.

**Figure 2:**
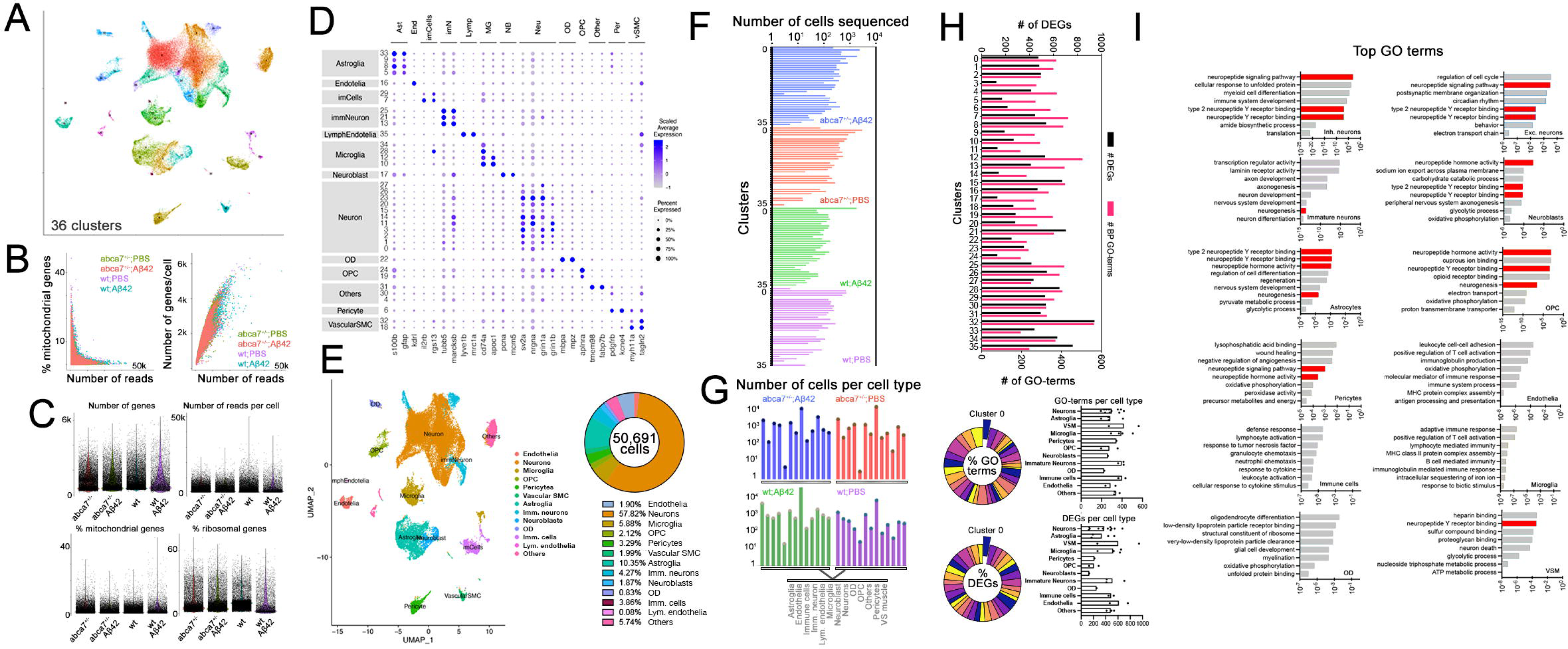
Single cell transcriptomics in *abca7* knockout zebrafish. (A) Combined UMAP plot for single cell transcriptomics from wild type + PBS, wild type + Aβ42, abca7^+/-^ +PBS and abca7^+/-^ + Aβ42 samples. 36 cell clusters identified. (B) Number of mitochondrial reads and number of genes per cell. (C) Number of genes, number of reads per cell, percentage of mitochondrial gene expression and percentage of ribosomal genes in every sample. (D) Dot plot for cell type markers identify distinct cell types. (E) UMAP plot classifying cell types and their percent prevalence in a total of 50,691 cells sequenced. (F) Distribution of the number of cells sequenced in all 36 clusters in every sample. (G) Number of sequenced cells per cell type. (H) Number of differentially expressed genes (DEGs, black) and gene ontology terms related to biological processes (magenta). Lower panels indicate pie chart of the percentage of GO-terms and DEGs per cluster, and the numbers of GO-terms and DEGs per cell type as bar graphs. (I) Tally for top GO terms in distinct cell types. neuropeptide signalling pathway and neurogenesis are enriched in neurons, neuroblasts, astroglia, oligodendrocyte progenitors and vascular smooth muscle cells.

To determine the differentially expressed genes, we used FindMarkers functions of Seurat and found the genes that are differentially expressed in *abca7*^+/-^ animals in comparison to wild types in the presence and absence of Aβ42 toxicity (*abca7*^+/-^ vs wild type, and *abca7*^+/-^/Aβ42 vs wild type/Aβ42) (Figure 2H, Supplementary Data 2). When we selected the top enriched GO-terms in every cell type, we observed that neuropeptide Y signaling and neurogenesis were typically represented in astroglia, excitatory and inhibitory neurons, oligodendrocyte precursor cells, and pericytes (Figure 2I). Interestingly, in oligodendrocytes (OD), these processes were not altered. In ODs, DEGs enriched lipid metabolism and lipoprotein activity-related processes, as shown before^40,41, 50^. These results suggested that Abca7 function was related to neuropeptide Y signaling. The regulatory role of Abca7 on astroglia could be a previously unidentified molecular process, that links Aβ42-induced astroglial proliferation and regenerative neurogenesis manifesting in zebrafish^7-9,22,24,26^ but not in mammals^19,20,23,29,51,52^. This neurogenic response could also be related to sustained synaptic integrity through enhanced resilience of neuronal networks.

To determine how the gene expression in every cell cluster in the individual experimental and control groups change with respect to Aβ42 and *abca7* knockout, we generated the UMAP plots for four groups: wt, *abca7*^+/-^, wt/Aβ42 and *abca7*^+/-^/Aβ42 (Figure 3A-C) and generated differential gene expression in these clusters (Supplementary Data 3). Although all cell clusters contained sequenced cells, neuronal cluster 0 was significantly more abundant in only wt/Aβ42-injected samples (green in Figure 3C, Supplementary Figure 2), compared to *abca7*^+/-^/Aβ42 cells (violet in Figure 3C, Supplementary Figure 2). This cluster was also present in phosphate buffer saline (PBS, control vehicle) injected wild type and *abca7*^+/-^ samples (Figure 3B), suggesting that Aβ42-injection might be inducing a specific set of neuronal mechanisms, reversable by *abca7* knockout. To determine the marker genes of every cell cluster, we plotted the expression level of the top-5 expressed genes in every cell cluster (Figure 3D, Supplementary Data 4) and found that neuropeptide y (*npy*) was the most expressed gene in Aβ42-related neuronal cluster 0. By plotting expression levels per cell type in violin plots (Figure 3E), we found that *npy* expression was significantly increased in many cell types upon Aβ42-injection but most prominently in neurons (Figure 3E, lower panel, blue), and *abca7*^+/-^ knockout reduced *npy* expression significantly (Figure 3E, F). Immunohistochemical staining for NPY confirmed the expression throughout the telencephalon (Figure 3G), which significantly reduced following Aβ42 injection in *abca7*^+/-^ knockout (Figure 3H). These results imply that amyloid-induced *npy* expression in neurons requires *abca7* activity.

**Figure 3:**
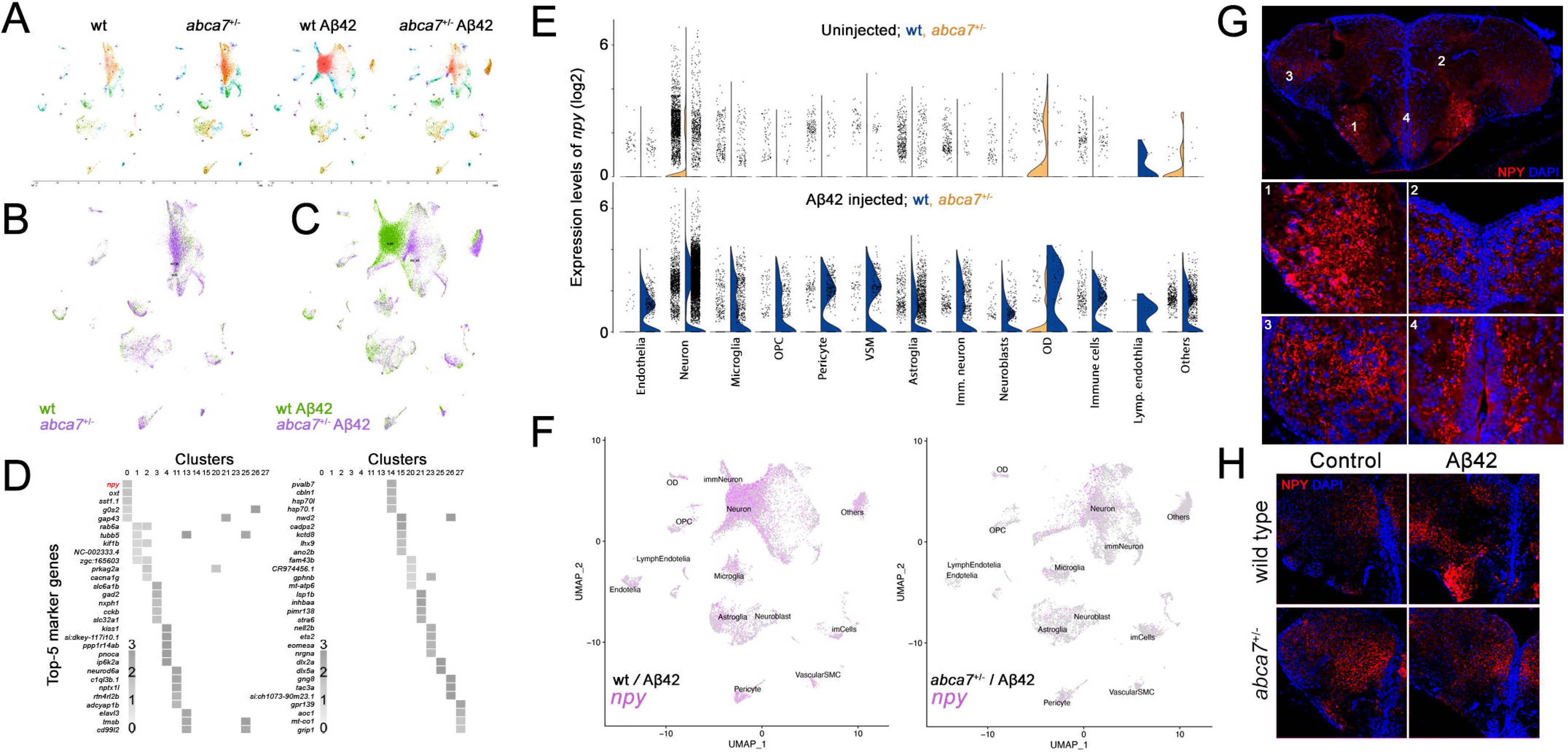
***npy* expression is reduced with *abca7* knockout.** (A) UMAP clustering for individual experimental groups of wild type + PBS, wild type + Aβ42, abca7^+/-^+PBS and abca7^+/-^ + Aβ42. (B) Combined UMAP for wild type (green) and *abca7* knockout (violet) shows overlapping cell clusters. (C) Combined UMAP for wild type + Aβ42 (green) and *abca7*^+/-^ + Aβ42 (violet) identifies a specific neuronal cluster (Cluster 0) that is enriched in Aβ42-treated wild type animals but this cluster is significantly diminished in abca7-knockout. (D) Top 5 marker genes for every cell cluster identifies neuropeptide y (*npy*) as the top marker for cluster 0. (E) Violin plots for *npy* expression. Upper plot is for wild type (blue) and *abca7* knockout (yellow) without Aβ42 injection. Lower plot is for wild type (blue) and *abca7* knockout (yellow) with Aβ42 injection. *npy* expression is significantly upregulated in a variety of cells, neurons being the highest upregulation. Abca7 knockout significantly reduces *npy* expression, mostly in neurons. (F) Individual expression plots for npy in wild type (left) and *abca7* knockout (right) with Aβ42 injection. Cluster 0, expressing *npy* is strongly reduced. (G) Immunofluorescence in control brains for Npy with DAPI counterstain. High magnification panels show indicated locations of the telencephalon. (H) Immunofluorescence for Npy with DAPI counterstain in wild type and *abca7* knockout animals with or without amyloid toxicity.

### Npy is required for neurogenesis and synaptic integrity

To predict the target cells of *npy*, which is primarily expressed in neurons and astroglia (Figure 4A), we determined the expression of neuropeptide receptors (Supplementary Figure 3). The most highly expressed receptor was *npy1r*, and its expression was predominantly in neurons (Figure 4A, Supplementary Figure 3), Since we observed a change in astroglia proliferation with *abca7* knockout after Aβ42 injection (Figure 1), we hypothesized that neuronal Npy signaling could have a direct effect on neurons and consequential effect on astroglia through a secondary downstream signalling pathway. When we compared the most enriched 10 GO terms in astroglia between *abca7*^+/-^/Aβ42 and wt/Aβ42 (Supplementary Data 2), we found a significant enrichment in neurogenesis and neuronal differentiation programs (Figure 4B). The differentially expressed genes in the top 10 GO terms included upregulated quiescence-related genes *id1*, *dla*, *notch1a*, as well as gliosis-related genes *s100b* and *gfap*, while several genes related to neurogenic potential and proliferation such as *her4.1, stmn2a, sparc*, and *hey1* were downregulated (Figure 4C). Interestingly, we also determined a downregulation in *ngfra*, which we have previously shown to be a key regulator of Aβ42-dependent astroglial proliferation and neurogenesis in adult zebrafish brain ^7,8^ and sufficient to induce proliferation and neurogenesis from otherwise non-neurogenic astroglia in APP/PS1dE9 mouse model of Alzheimer’s disease ^26^. When we detected the *ngfra* expression in individual cell types in comparison of *abca7*^+/-^/Aβ42 to wt/Aβ42, we found that *ngfra* expression was significantly reduced in astroglia and immature neurons (Figure 4D, E), suggesting that *abca7*-dependent regulation of astroglial proliferation could be via *ngfra*, a determinant of neurogenic potential of astroglia ^8,26^. We previously showed that Bdnf is an intermediate for the neuro-glial crosstalk for regulation of astroglia proliferation^8^, and in this study we found that *bdnf* expression overlaps with *npy* expression in zebrafish brain (Supplementary Figure 4). Therefore, we hypothesized that the *npy*-expressing neurons and *ngfra*-expressing astroglia could be communicating through Bdnf. To test this, we determined the *bdnf* expression and changes in our dataset. We found that *bdnf* expression was significantly reduced in *abca7*^+/-^ neurons in comparison to wild type neurons under Aβ42 toxicity (Figure 4F), suggesting that alterations in astroglial proliferation and neurogenicity in *abca7*^+/-^ animals could be due to reduced *bdnf* in neurons and reduced *ngfra* in astroglia.

**Figure 4:**
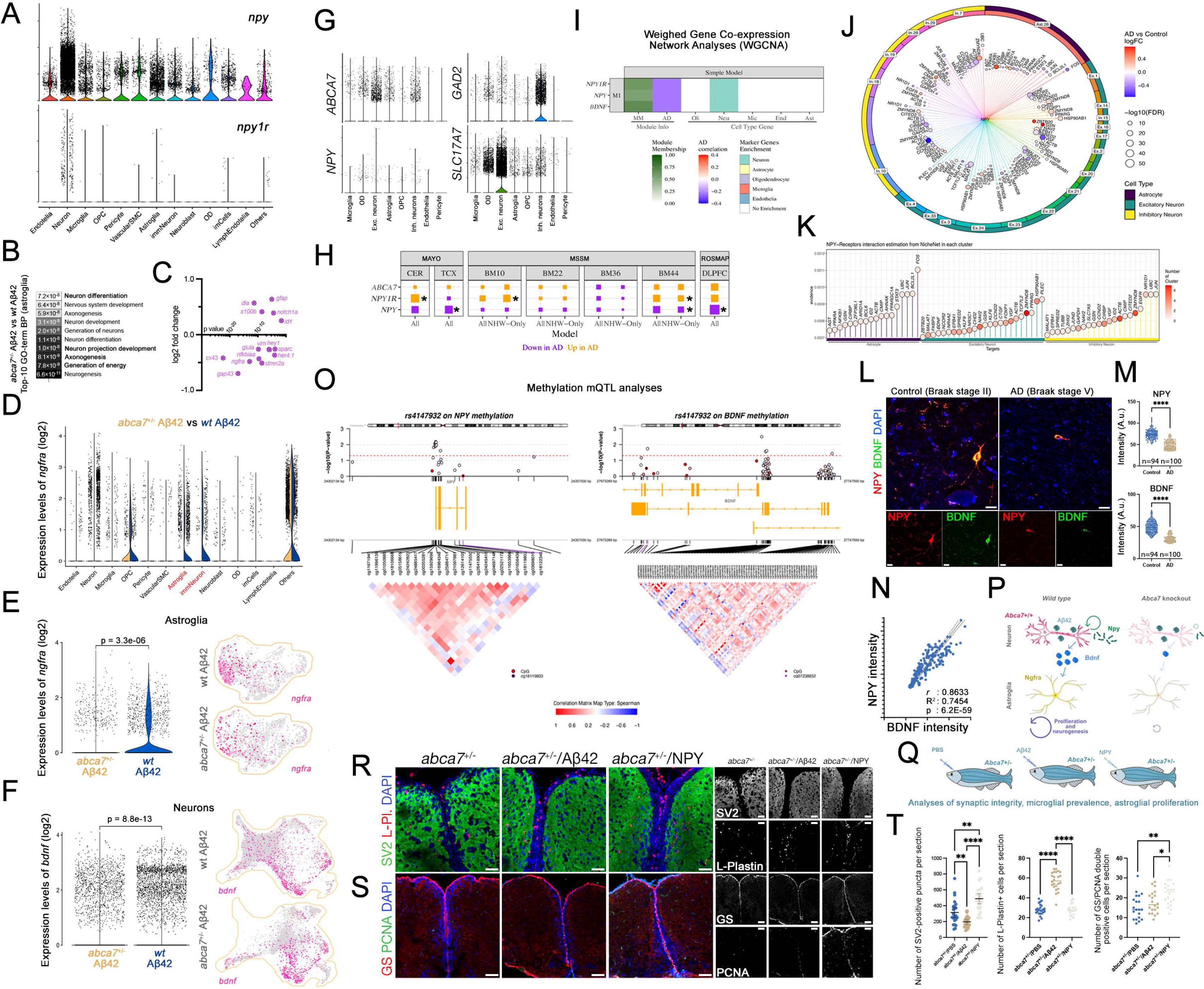
***npy* is required for maintaining synaptic integrity and astroglial proliferation.** (A) Violin plot for *npy* and *npy1r* expression in every identified cell type in zebrafish. (B) Top 10 enriched GO terms when *abca7*^+/-^ + Aβ42 and wild type + Aβ42 were compared indicate altered neurogenesis mechanisms in astroglia. (C) Neurogenic genes are downregulated; quiescence genes are upregulated in astroglia after *abca7* knockout. (D) Violin plot for expression of *ngfra* in astroglia when *abca7*^+/-^ + Aβ42 is compared to wt *+* Aβ42. (E) Expression of *ngfra* is strongly reduced in astroglia. (F) Ngfra ligand Bdnf is significantly reduced in neurons. (G) Expression of *ABCA7* and *NPY* in human brains (data reanalyzed from ^55^), showing co-expression in neurons. (H) Expression analyses in various AMP AD brain transcriptome datasets indicate uniform downregulation of *NPY* with AD. Asterisks indicate statistical significance (p < 0.05) (I) Weighed gene co-expression network analyses based on Mayo Clinic temporal cortex (TCX) RNAseq data indicate *NPY, NPY1R* and *BDNF* expressions are negatively correlating with AD and are associated with neurons. (J) *NPY*-centered NicheNet analyses in human brain datasets of single nucleus RNA sequencing. Excitatory neurons cluster 1showed reduced *BDNF* in AD in an *NPY*-dependent manner. (K) NPY-receptor interaction estimation and altered gene expression in astrocytes, excitatory neurons, and inhibitory neurons. (L) Double immunohistochemistry for NPY and BDNF in postmortem control and AD brains. (M) Quantification results of NPY and BDNF intensities in neurons. Asterisks denote p<1.0E-15. (N) Correlation analyses between BDNF and NPY in 194 neurons analyzed. (O) CpG methylation mQTL analyses with *ABCA7* variants in humans. The epigenetic changes on *NPY* and *BDNF* are shown. Red line indicates statistical significance cutoff. Every circle represents one methylation site. Blue indicates hypermethylation, red indicates hypometylation. Yellow bars indicate the gene location and coding direction. Enlarged region lists all methylation sites in the corresponding genomics window. Correlation matrix between different methylation sites are depicted as positive (red) and negative (blue). *ABCA7* variants exert methylation changes in *NPY* and *BDNF* promoters in a highly correlated manner. (P) Working model for Abca7-dependent Npy-mediated regulation of astroglial proliferation. (Q) Experimental setup for investigating the effect of NPY in mediating amyloid-induced phenotypes in synaptic integrity, microglial numbers and astroglial proliferation. (R) Immunofluorescence for SV2 and L-Plastin with DAPI counterstain in *abca7*^+/-^, *abca7*^+/-^ + Aβ42 and *abca7*^+/-^ + NPY samples. Black and white panels indicate individual fluorescent channels. (S) Immunofluorescence for GS and PCNA with DAPI counterstain in *abca7*^+/-^, *abca7*^+/-^ + Aβ42 and *abca7*^+/-^ + NPY samples. Black and white panels indicate individual fluorescent channels. (T) Quantification for number of SV2-positive synaptic puncta, L-Plastin-positive microglia, and GS/PCNA double-positive proliferating astroglia. n = 4 animals used for every group. Raw data: Supplementary Data 1.

### NPY-BDNF-NGFR axis is associated with Alzheimer’s disease

While brains in zebrafish and humans are evolutionarily distinct, pathological mechanisms of AD may be parallel ^6,7,14,15,17-21,26,28,29,51^, exemplified by the remarkable similarity of molecular changes in neurons after amyloid toxicity ^51^. Thus, we hypothesized that NPY and BDNF signaling might be altered in AD patients. To test this, we first determined the expression of *ABCA7* and *NPY* in single nuclei transcriptomics study of human brains ^53-55^ and found that *ABCA7* and *NPY* were expressed in inhibitory and excitatory neurons, while *ABCA7* expression was also present in microglia and oligodendrocytes (Figure 4G, Supplementary Data 5). We analyzed human bulk RNA sequencing in brain tissue from the Mayo Clinic, ROSMAP and Mt Sinai Brain Bank. In cerebellum (CER) and superior temporal gyrus (TCX) regions ^56,57^, Brodmann areas BM10, BM22, BM36 and BM44 brain regions ^58^ dorsolateral prefrontal cortex (DLPFC)^59^, we found that *NPY* had a consistent downregulation trend in all cohorts (significantly downregulated in AD in Mayo-TCX, ROSMAP and MSBB BM44; p < 0.05; Supplementary Data 5, Supplementary Table 2). NPY receptor *NPY1R* is upregulated in Mayo-CER, MSBB BM10 and BM44 (p < 0.05, Figure 4H, Supplementary Data 5). In Mayo-CER and Mayo-TCX, *BDNF* is significantly downregulated in AD in (FDR = 0.015 and 2.1E-6, respectively; Supplementary Data 5).

To determine whether *NPY* and *NPY1R* were associated with AD through co-expression modules of gene expression, we performed WGCNA analyses on the Mayo-TCX brain samples and found that this module was significantly downregulated in AD and enriched with neuronal genes (Figure 4I). Through NicheNet ^60^ analyses, which predicts the cellular interactions through candidate ligand-receptor pairs, we found that in humans, *NPY* interacted with several cell types to regulate the expression of various genes in target cells (Figure 4J, Supplementary Data 6). Interestingly, in the largest excitatory neuron cluster, the highest gene expression change was observed in *BDNF* (Figure 4J, K), suggesting that NPY-BDNF-NGFR axis was similar in humans as in zebrafish. When we compared the NicheNet targets in humans (Figure 4J) to our previous dataset in mouse model of AD where the molecular regulation of induced NGFR signalling was analyzed ^26^, we found that 37% (18/49) of the genes that were potential NPY targets in humans (Figure 4J, K) were also differentially regulated after *NGFR* expression in the hippocampus of the APP/PS1 mouse model of AD (Supplementary Figure 5). This finding suggested potential crosstalk mechanisms between NPY-expressing neurons and NGFR-expressing astroglia via BDNF. We confirmed this interaction by immunohistochemical staining in postmortem human brains (controls vs AD), and found that in AD brains, NPY and BDNF colocalize in neurons, and their levels were reduced in AD compared to controls (Figure 4L, M). NPY and BDNF levels in the brain significantly correlated in neurons (R^2^: 0.74, p: 6.2E-59, Figure 4N). The reduction of Bdnf with amyloid was also consistent in the *abca7* knockout zebrafish model (Supplementary Figure 6), similar to Npy (Figure 3H).

### Loss-of-function variants in *ABCA7, NGFR* and *BDNF* segregate in families with AD

Our findings propose that *ABCA7*-dependent NPY activity could establish a neuro-glial crosstalk through BDNF and NGFR that regulates astroglial physiology, and genetic variants in these genes may be associated with familial AD. Results from family-based studies (demographics: Supplementary Table 3) showed that *ABCA7* gene was completely (Supplementary Table 3, Supplementary Figure 7) and incompletely (Supplementary Table 4) segregating in both Hispanic and the white, non-Hispanic AD families. Two variants from the *ABCA7* genes were segregating in the Hispanic and two variants were segregating in the white, non-Hispanic AD families (Supplementary Table 4). This indicates the strong association of *ABCA7* with familial AD. Additionally, *BDNF* gene was completely segregating (Supplementary Table 3 and 4) in the white, non-Hispanic families but not segregating in the Hispanic families. In the *BDNF* gene, one variant was segregated in white, non-Hispanic families but co-segregated with *APOE* ε4 (Supplementary Table 4). Taken together, *ABCA7-NPY-BDNF-NGFR* axis appears relevant to familial AD and likely more pronounced in the white, non-Hispanic ancestry.

### *ABCA7, NPY* and *NGFR* co-expression is associated with higher Braak staging in human brains

*NPY* expression reduced in AD (Figure 4, Supplementary Table 2, Supplementary Data 5), thus we investigated whether Braak stages were also associated with co-expression among our five genes (*ABCA7, BDNF, NPY, NPY1R, NGFR*), adjusting for sex, age at death, RIN, and race (Supplementary Table 5). Indeed, our data confirmed that two statistical interactions were significant after multiple testing correction: *NPY*NGFR* (OR= 2.06 [1.28-3.70]) and *ABCA7*NGFR* (OR= 3.29 [1.48-8.72]) consistent with our previous hypothesis that the *ABCA7-NPY* axis can act through *NGFR* signalling and this interaction might be associated with increasing Braak stages in AD. By depicting co-expression patterns of *ABCA7, NPY* and *NGFR*, we found that *NGFR* expression inversely correlates with *ABCA7* and *NPY* expression only when AD pathology is absent or mild (BRAAK 0-4), while in AD pathology (BRAAK 5-6), this correlation was lost (Supplementary Figure 8). These results suggest that in mild cases or controls, reduced *ABCA7* or *NPY* leads to an increase in *NGFR* as a potential protective reaction, which can be absent in AD, when *NPY* and *ABCA7* expression reduces, AD brains cannot elevate *NGFR* levels. This statistically significant differential interaction dependent on the Braak stage was consistent with our previous study where activating *NGFR* in the brains of APP/PS1dE9 mice reduced amyloid and Tau burden^26^.

### *ABCA7* variants are associated with epigenetic regulation of *NPY, BDNF* and *NGFR*

To further test the potential regulatory role of *ABCA7* on *NPY*, *BDNF* and *NGFR*, we identified potential mQTL signals of *ABCA7* variants in humans using New York Brain Bank and NIA AD-FBS/NCRAD cohorts^61,62^ (Figure 4O). According to the regional plots (Figure 4O), we observed that the genetic variants at *ABCA7* have nominally significant mQTL effects (P<0.05) on the methylation at loci of *ABCA7, NPY, BDNF,* and *NGFR* at transcriptional regulatory regions (Figure 4O, Supplementary Data 7). This indicated that *ABCA7* might influence the expression of *NPY*, *BDNF* and *NGFR* through epigenetic regulation and potentially through transcriptional activity.

### Ectopic NPY rescues the amyloid-induced synaptic degeneration and impaired astroglial proliferation in *abca7* knockout

Considering the human data, we hypothesized that *abca7*-expressing neurons could express *npy* in amyloid toxicity conditions as a stress response, which could potentiate autocrine signalling through *npy* receptors and allow production of Bdnf that regulates proliferation in astroglia through Ngfra in zebrafish (Figure 4P). This cascade could be impaired by *abca7* knockout through reducing *npy* availability, *bdnf* expression, *ngfra* signalling, and in turn modulating the synaptic integrity and astroglial proliferation. To test this hypothesis, we injected human NPY to *abca7^+/-^* zebrafish treated with Aβ42 and compared this with non-injected group (Figure 4Q) for investigating whether NPY rescued the alterations in astroglial proliferation, synaptic degeneration, and microglial activity. By performing immunohistochemical stains for synaptic vesicle protein SV2 as a neuronal marker and L-Plastin as a microglial marker (Figure 4R), and GS as an astroglial marker and PCNA as a proliferating cell marker (Figure 4S), we found that NPY was sufficient to rescue the reduction of SV2-positive synaptic puncta in Aβ42-treated *abca7*^+/-^ neurons and increase the astroglial proliferation, while not affecting the number of microglial cells (Figure 4T). We also found that in zebrafish, NPY injection can restore the amyloid induced BDNF expression that is abrogated by *abca7* knockout (Supplementary Figure 6). Combined with the effects of *Abca7* knockout on these outcomes (Figure 1), our results suggested that Abca7-dependent Npy activity in zebrafish regulates synaptic integrity and glial proliferation while Abca7 was required for microglial proliferation independent of Npy.

## Discussion

In this study, we generated the first stable adult *abca7* gene knockout in zebrafish and investigated the impact of heterozygous *abca7* deletion in zebrafish brains in the presence and absence of amyloid toxicity. The *abca7*^+/-^ zebrafish exhibited altered responses to Aβ42 toxicity, with reduced astroglial proliferation and synaptic density. Single cell transcriptomics revealed significant changes in gene expression of *abca7* loss-of-function variants in the amyloid toxicity context. Neuropeptide Y (*npy*) levels increased in neurons due to Aβ42 in wild type zebrafish but decreased with *abca7* knockout, potentially abrogating a beneficial response in neurons. Injecting human NPY restored altered processes and BDNF expression in *abca7*^+/-^ zebrafish, suggesting a therapeutic potential. Comparative human studies validated the changes in *NPY* expression in human AD brains, and its regulatory signaling through *BDNF*. Our results uncovered a potential role of *abca7* as a neural resilience and neurogenic factor.

This study confirms and augments the role of *abca7* in multiple cellular mechanisms including neuronal response to amyloid, immune system activity, and astroglial proliferation through potential involvement of Neuropeptide Y, a 36-aminoacid peptide with neuroprotective effects and anti-inflammatory roles ^63-71^. Among the highest expression of NPY in human brain is in hippocampus ^72-74^, the primary neurogenic location. *NPY* expression was reduced in AD in humans ^63,73, 75, 76^ and animal models ^68,69^, which indicated a protective and beneficial effects of this gene for the homeostatic health of the brain. A recent large single nucleus transcriptomics study in human brains also confirms our findings. This study identified *NPY* and *SST* expressing inhibitory neurons reduce *NPY* expression in AD, and suggested that such alterations could constitute a mechanism that impairs the brain resilience with aging and AD ^77^. Our study also identified *NPY* and *SST* to be co-expressed in inhibitory neurons (Supplementary Figure 9), and *NPY* decreased in AD in association with Braak staging. We add further mechanistic insight into the role of NPY-expressing neurons in brain resilience against AD and propose that this mechanism is *ABCA7*-dependent. Our transcriptomics and functional studies suggest that NPY might be related to sustained neurotrophic effect through regulation of the availability of BDNF. Our NicheNet analyses also supported change in NPY in human brains was linked to reduced BDNF in neurons in human brains (Figure 4). These findings were consistent with the previous observations that Npy positively regulates *Bdnf* expression in rat cortical neurons exposed to Aβ42 ^67^.

We also found that the *abca7* knockout reduced the proliferation of astroglia, which was consistent with the reduced expression of *ngfra*, a key regulator of astroglial proliferation and neurogenesis in zebrafish brain after amyloid toxicity. Astroglia, as neural stem cells, bear the potential as a regenerative therapy agent for a variety of medical conditions ^52,78-83^. The use of endogenous NSCs may limit tissue loss and improve tissue integrity, with potential therapeutic applications in neurodegeneration ^23,84^. Thus, a plausible approach could be inducing endogenous astroglia to generate newborn neurons and replace the lost ones, thereby increasing brain resilience to neurodegeneration. Previous studies showed that induced *in vivo* expression of *Npy* in mouse model of AD can increase the neural progenitor cell proliferation and neurogenesis ^65^. We found that *abca7*-dependent consistently reduced *npy* expression which correlated with reduced proliferative response of astroglia after amyloid toxicity, and this reduction could be rescued by ventricular injection of human NPY.

Taken together, we propose that ABCA7 is required for increased NPY as a response to AD related neuropathology, and NPY activity is crucial for synaptic integrity and astroglial proliferation in AD. These two processes may help to maintain a homeostatic brain resilience, which might be impaired in AD. This hypothesis is supported by our genetic association studies where *ABCA7* and *NPY* expression is associated with *NGFR* in low Braak stages of AD but not in high Braak stages (Supplementary Figure 8). In conjunction with our previous findings that NGFR imposes proliferative and neuroregenerative potential to the astroglia of APP/PS1dE9 AD mouse model and reduces the amyloid and Tau burden, these results support the protective role of NGFR signalling in mammalian brains against AD is impaired with advancing AD pathology. In AD brains, reduction in *NPY* expression was present in all regions but prominently in the hippocampal region suggesting that *NPY* signaling could regulate neurogenesis. The hippocampus, one of the first brain regions to be affected in AD, contains neural progenitor cells (NPCs) that continue to generate new neurons for the course of adult hippocampal neurogenesis ^85,86^, which is impaired in AD ^87,88^. Blocking neurogenesis exacerbates neuronal loss and cognitive decline, while neurogenesis in combination with BDNF treatment ameliorates the behavioral readouts ^89^. Therefore, neurogenesis together with neurotrophic mechanisms can sustain the resilience of the brain and ability to cope with neurodegeneration. Our results suggest that *abca7*, through regulating both neurogenesis and neurotrophic factor expression, may be necessary for protection or resilience against AD neuropathology, also consistent with AD risk associated with both common and rare *ABCA7* genetic variants. Our results propose a new biological crosstalk between *ABCA7*, *NPY*, *BDNF,* and *NGFR* which are genetically associated with AD ^5^.

Although we have established a link between *abca7* and *npy*, and subsequent effects on neural resilience and astroglial proliferation, the cellular mechanisms through which *abca7* loss-of-function alters *npy* expression, requires further investigation. ABCA7’s impact on lipid metabolism is believed to influence Aβ processing, potentially leading to the aggregation of Aβ neuritic plaques, a hallmark of AD. Moreover, ABCA7’s interactions with other proteins involved in lipid metabolism and immune response regulation further underscore its role in AD pathology, where NPY could partake. A possible mechanism could involve the structural and functional assembly of lipid rafts through ABCA7 activity, as it has been demonstrated in lymphocytes ^42^ and hypothesized as a general regulatory pathway before ^90^. This is consistent with the proposed role of ABCA7 in regulating lipid metabolism and localization on the cell membrane ^40,41, 43^. NPY also interacts with the lipid bilayer on cell surface in a conformation-dependent manner ^91^, and altered lipid content and organization could affect the activity of NPY directly or indirectly through its various receptors such *NPY1R*, which we found to be expressed in neurons in humans and zebrafish. Previous studies demonstrated that NPY and BDNF can stimulate each other’s expression in a feed-forward loop ^67,69, 70^, suggesting that loss of *ABCA7* function could impair this neurotrophic signaling by disrupting the lipid-mediated NPY signalling, resulting in reduced BDNF expression in neurons. This downregulation of BDNF might affect astroglial behavior, potentially influencing processes such as proliferation and neurogenesis through NGFR-mediated pathways. This crosstalk highlights the interconnectedness of lipid metabolism, NPY signaling, and BDNF-mediated effects on cellular responses in an *ABCA7*-dependent mechanism and suggests novel therapeutic targets and directions.

We also note the limitations of our study. We identified expression of *npyr8a* in zebrafish, which can potentially bind to Npy, yet the functional role of this receptor subtype needs further investigation. Although *Abca7* knockout changes microglial activity and numbers, NPY does not, suggesting that Abca7’s role on microglial dynamics is NPY-independent and requires further analyses. ABCA7 is a critical regulator of lipid metabolism, and it could regulate secretion of neuropeptides or functional maturation and organization of receptor complexes on lipid bilayer of the membrane. This potential mechanistic relationship requires further analyses on lipid content of Abca7 knockout animals and can point towards a general alteration in secretory pathways. Further studies should also focus on the potential protective role of *ABCA7* in the context of other neurodegenerative pathologies, given that *ABCA7* variants are also associated with risk of non-AD neurodegenerative diseases ^33,36^.

## Materials and Methods

### Ethics statement and clinical samples

All animal experiments were performed in accordance with the applicable regulations and approved by the Institutional Animal Care and Use Committee (IACUC) at Columbia University (protocol number AC-AABN3554). In every experimental set, animals from the same fish clutch were randomly distributed for each experimental condition. Animals were handled with caution to reduce suffering and overall animal numbers. Human brain samples were obtained from New York Brain Bank within institutional regulations of Columbia University. The clinical data analyses conducted at Columbia University and Mayo Clinic were approved by the appropriate Institutional Review Boards.

### Gene editing and generation of *abca7* knockout zebrafish line

Cas9 mRNA synthesis: The template DNA containing the capped Cas9-mRNA, (pCS2-Cas9) was linearized by *NotI* digestion and then purified by reaction purification kit (Invitrogen). Using mMESSAGE mMACHINE SP6 kit (Invitrogen), capped Cas9-mRNA was synthesized and then purified by ethanol precipitation as indicated in the manufacturer protocol. sgRNAs targeting middle (exon13) and end region (exon46) of the *abca7* gene were selected through the CRISPRscan (https://www.crisprscan.org) web tool. Oligo for middle region sgRNA: 5-taatacgactcactataGGATCATGGGTCTGGGCTCAgttttagagctagaa-3 and oligo for end region sgRNA: 5-taatacgactcactataGGGCAAATCAAAAAGTCCATgttttagagctagaa-3 were prepared based on a cloning-free method. Each sgRNA oligo was used as forward primer in a a PCR reaction with gRNA universal tail primer 5-AAAAGCACCGACTCGGTGCCACTTTTTCAAGTT GATAACGGACTA-3 as reverse primer. PCR reaction for each sgRNA was set up with a 100 µl reaction volume containing 1 X Q5 buffer, 2 µM each primer, 0.2 mM each dNTP, 2 µl Q5 DNA Polymerase (NEB). Amplification was done with an initial denaturation at 95 °C for 30 sec followed by 40 cycles at 95 °C for 10 sec, 60 °C for 10 sec, and 72 °C for 10 sec, and a final extension at 72 °C for 5 min. The PCR products were then purified by PCR purification kit (Invitrogen) and used as template DNA to generate each sgRNA by in vitro transcription using MEGAshortscript T7 kit (Invitrogen). The size and quality of the gRNAs were visualized in %3 agarose gel electrophoresis. The mix of sgRNAs and capped mRNA was injected directly into one-cell stage zebrafish embryos. For each embryo, 2 nl solution containing 600 pg Cas9 mRNA and 100 pg sgRNA from each was injected. To confirm the deletion, specific primers binding on the upstream and downstream regions of the gRNA target sites were designed. 10 single 2-day post injection (dpi) embryos were collected individually and each of them was put into a 1.5 ml Eppendorf tube. Genomic DNAs of single embryos were prepared by alkaline lysis method^92^. . 3 µl genomic DNA solution from each embryo was used in a 25 µl PCR reaction containing 1X Taq buffer, 2 µM each primer, 0.2 mM each dNTP and 0.5 µl Taq DNA polymerase. PCR fragment indicating the deletion was visualized in %2 agarose gel electrophoresis. Subsequent DNA sequencing confirmed the partial deletion in abca7 gene. Injected embryos were raised and then 4-month chimeric F0 candidates were crossed to the WT AB fish and their 2-day embryos were screened by PCR and genotyped by Sanger sequencing. Potentially positive F1 embryos were raised to adulthood (4-month) and then were screened by Sanger sequencing following the PCR on the target site of the genomic DNA prepared from their tail fins. In the end, 16 F2 fish having the partial deleted *abca7* gene were identified.

### Cerebroventricular microinjection, tissue preparation and immunohistochemistry

Cerebroventricular microinjections (CVMI) into adult zebrafish brain were performed as described^13,24^. 8-12 months old *abca7*^-/+^ heterozygous knockouts and their respective age-matched wild type siblings of both sexes were used for the experiments. The animals were injected with the followings: PBS (control), human amyloid-beta42 (Aβ42, 20 µM), human Neuropeptide-Y (10 μg/ml). Total injection volume was 0.5 – 1 μl. At 3 days post injection (dpi), animals were euthanized and subjected to histological tissue preparation. The zebrafish heads were dissected and fixed overnight at 4°C using 4% paraformaldehyde. After several washes the heads were incubated overnight in 20% Sucrose with 20% ethylenediaminetetraacetic acid (EDTA) solution at 4°C for cryoprotection and decalcification. The following day, fish heads were embedded in cryoprotectant sectioning resin O.C.T. and cryosectioned into 12-μm thick sections on SuperFrost Plus glass slides. For immunohistochemistry, the sections were dried at room temperature, followed by washing steps in PBS with 0.03% Triton X-100 (PBSTx). Primary antibodies were applied overnight at 4°C. Next day, the slides were washed 3 times with PBSTx and then appropriate secondary antibodies were applied for 2 hours at room temperature. The slides were then washed several times before mounting using 70% glycerol in PBS. Antibodies used are listed in Supplementary Table 1. For antigen retrieval of PCNA and SV2, slides were heated in 10 mM Sodium acetate at 85°C for 15 minutes before primary antibody incubation. For antigen retrieval of NPY, slides were heated in 50 mM Tris-HCl at 95°C for 5 minutes before primary antibody incubation.

### Imaging, quantifications, and statistical analyses

Images were acquired using a Zeiss confocal LSM800 microscope in a 20X or 40X objectives with tiles/z-stack function wherever necessary. 6 telencephalon sections between the caudal end of the olfactory bulb and anterior commissure were used per animal for the quantitative analyses, with at least 4 animals per experimental group (n=4) in each of the experimental setup. The quantification of SV2-positive synapses was performed using 3D object counter module of ImageJ software with a same standard cut-off threshold for every images. For quantification of microglia, the number of cells immunoreactive to L-Plastin were counted. For quantification of PCNA+GS double-positive cells, overlap of PCNA-positive nuclei with GS-positive cell were counted. The statistical analyses were performed using GraphPad Prism (Version 9.5.1) for one-way ANOVA followed by a Sidak’s multiple comparison test or Dunnett’s multiple comparison test wherever applicable. Error bars shown are the s.e.m. and asterisks indicate significance according to: *: p<0.05, **: p<0.01, ***: p<0.001. p>0.05 is considered not significant (n.s.).

### Single cell sequencing in zebrafish and data analyses

The telencephalon of the 10-month-old fish were dissected in ice-cold PBS and directly dissociated with Neural Tissue Dissociation Kit (Miltenyi Biotec, Cat. No. 130-092-628) as described previously^24^. After dissociation, cells were filtered through 40 μM cell strainer into 10 mL 2% BSA in PBS, centrifuged at 300 g for 10 min, and resuspended in 4% BSA in PBS. Viability indicator dyes Sytox Blue (Invitrogen, Cat No. S34857) and Dycle Ruby (Invitrogen, Cat. No. V10309) were used to sort the cells by FACS. Samples from abca7 heterozygous knock-out and WT-negative siblings were sorted in separate FACS machines (Sony MA900-FP) to ensure as minimum incubation time for the samples placed on ice (Supplementary Data 8). The resulting single cell suspension was promptly loaded on the 10X Chromium system^93^. 10X libraries were prepared as per the manufacturer’s instructions. Generated libraries were sequenced via Illumina NovaSeq 6000 as described^8,9,26,51,94-96^. In total, 54,482 cells were sequenced. The raw sequencing data was processed by the Cell Ranger Single Cell Software Suite (10X Genomics, v6.1.2) with the default options. On average, 96.65% of the total 1,70 billion gene reads were aligned to the zebrafish genome release GRCz11 (release 105). The resulting matrices were used as input for downstream data analysis by Seurat^97^.

### Read alignment, and quality control

The single cell expression matrices were read by Read10X function of Seurat^98^ (version 4.1.3) R package. The Seurat objects were created by filtering out the any cells with less than 200 expressed genes, and with genes expressed in less than 3 cells. After filtering out the low-quality cells, seurat objects were normalized, and the top 2,000 variable genes were used for further analyses. After identifying the anchors (FindIntegrationAnchors), the datasets were integrated (IntegrateData). We used DoubletFinder^99^ to identify and remove doublets, and the rest of the analyses were done on singlets. The integrated Seurat object included 50,691 cells with 26,122 genes. The data were scaled using all genes, and 30 PCAs (RunPCA) were identified. Cell clustering, marker gene analyses, differential gene expression and preparation of feature plots were performed as described^9,19,51,97,100^. Resolution of was used to identify the clusters. In total, 35 clusters were identified. The main cell types were identified by using *s100b* and *gfap* for Astroglia; *sv2a, nrgna, grin1a, grin1b* for Neuron; *pdgfrb* and *kcne4* for Pericyte; *cd74a* and *apoc1* for Microglia; *mbpa* and *mpz* for Oligodendrocyte; *aplnra* for OPC; *myh11a* and *tagln2* for vascular smooth muscle cells, *lyve1b* for lymphatic endotelial cells and *kdrl* for vascular cells^8,9, 26^. To find differentially expressed genes (DEGs), we used FindMarkers function of Seurat with 0.25 logfc.threshold. And. for GoTerm analyses, DEGs were used as described before^9^. The dataset can be accessed at NCBI’s Gene Expression Omnibus (GEO) with reasonable inquiries to the corresponding author.

### Expression analyses in AD cohorts

Analysis of the human AD bulk RNAseq data has been reported previously^26^. Briefly, bulk brain gene expression data was obtained the Accelerating Medicines Partnership - Alzheimer’s Disease (AMP-AD) datasets, available via the AD Knowledge Portal (https://adknowledgeportal.synapse.org). To assess differentially expressed genes (DEG) between AD and control, multiple linear regression models were used to compare the conditional-quantile-normalized expression values between AD and control participants while adjusting for covariates including RNA integrity number (RIN), sex, age at death, sequencing flowcell or batch, tissue source, and and participant race, where applicable, followed by multiple testing corrections using false discovery rate. For expression data from the Mount Sinai Brain Bank, AD vs control DEG was also performed in a subset of non-Hispanic white (NHW) participants. The associations between gene expression and Braak stage were assessed in TCX, CER, and DLPFC, following similar multiple linear regression models where age at death, sex, RIN, sequencing flowcell or batch, and APOE E4 allele dosage were adjusted. Multiple testing was corrected using false discovery rate. Cell-intrinsic DEG results were retrieved from previously published work^101^. The construction of the weighted gene co-expression network analysis (WGCNA)^102^ was described previously^103^. We reported the co-expression modules containing the genes of interest.

The single-nucleus RNAseq data^104^ were generated using frozen post-mortem tissue from the temporal cortex of 12 individuals with pathologically confirmed AD^105^ cases, and 12 age-, sex-matched control individuals with approval from the Mayo Clinic Institutional Review Board and written informed consent from the participants or their qualified next-of-kin. Briefly, to assess tissue quality, RIN was measured using RNA Pico Chip Assay (Agilent Biotechnologies, 5067-1513) on an Agilent 2100 Bioanalyzer using total RNA extracted from ∼20mg of brain tissue. Single nuclei suspension was collected on high-quality tissues (RIN > 5.5) using an established protocol with modification^106^. The nuclei were incubated with mouse anti-Human Nuclear Antigen [235-1] (Abcam, ab191181) antibody at 1:200 dilution and sorted using a BD FACSAria II sorter. Single cell RNAseq libraries were prepared using the Chromium Single Cell 3’ Gel Bead and Library Kit v3 (10X Genomics, 120237) and the Chromium i7 Multiplex Kit (10X Genomics, 120262) according to the manufacturer’s instructions. DNA libraries were sequenced at the Mayo Clinic Genome Analysis Core (GAC) using the Illumina HiSeq4000 sequencer.

Raw reads were aligned to human genome build GRCh38 and a premature mRNA reference file. After QC, log-normalization, and clustering of the snRNAseq data, cluster marker genes were calculated as the genes that are 1) expressed in over 20% AD and control nuclei, 2) overexpressed in the cluster (logFC > 0.25), and 3) the Bonferroni-adjusted p-value for the overexpression is less than 0.05 based on rank sum test. The cell types were subsequently assigned based on information from two approaches. First, the statistical enrichment of cluster marker genes in a list of marker genes^107^ for neurons, astrocytes, oligodendrocytes, endothelial cells, and oligodendrocyte progenitor cells (OPCs) was calculated using hypergeometric tests. Concurrently, the overlap of cluster maker genes and canonical cell marker genes for neurons (*SYT1*, *SNAP25*, *GRIN1*), excitatory neurons (*SLC17A7*, *NRGN*), inhibitory neurons (*GAD1*, *GAD2*), oligodendrocyte (*MBP*, *MOBP*, *PLP1*), astrocyte (*AQP4*, *GFAP*), microglia (*C3*, *CSF1R*, *CD74*), OPCs (*VCAN*, *PDGFRA*, *CSPG4*), endothelial cells (*FLT1*, *CLDN5*), and pericytes (*PDGFRB*) were assessed to assign the cluster cell type.

### *In silico* interaction mapping

NicheNet^108^ analysis tool was applied, in a targeted fashion, to study the interaction between *NPY* and genes in each cluster through NicheNet R package. Prior knowledge of ligand-target interaction has been compiled and optimized by NicheNet from multiple data sources to give a prior model which contains the regulation strength of ligands towards target genes. In this study, we assumed that *NPY* was the ligand based on previous works (references). Further, we required the NPY receptors being the following *NPY1R*, *NPY2R*, *CXCR4*, *NPY4R*, *NPY5R*, or *NPY6R*. For a given cluster that had NPY receptors expressed in it, the predicted target genes of NPY were those that 1) DEGs between AD and control cells, 2) expressed in >= 20% cells of the cluster, 3) among the top regulated genes of NPY according to the NicheNet prior model.

### Whole Genome Sequencing (WGS)

We analyzed WGS in 2,535 individuals from 522 families in EFIGA^62^ and AD-FBS cohorts. The NIA AD-FBS is the largest collection of familial AD worldwide^61^. The sequencing was performed at the New York Genome Center (NYGC) as described^109^, using one microgram of DNA, an Illumina PCR-free library protocol, and sequencing on the Illumina HiSeq platform. Variants were called using NYGC automated analysis pipeline which is based on CCDG and TOPMed recommended best practices^110^. Variant filtration was performed using Variant Quality Score Recalibration (VQSR at tranche 99.6%), sample missingness (>2%), depth of coverage (DP<10) and genotype quality (GQ>20). We then annotated high quality variants using ANNOVAR for population level frequency using Genome Aggregation Database (gnomAD), in-silico function using Variant Effect Predictor (VEP) and variant conservation using Combined Annotation Dependent Depletion score (CADD).

### Rare variant family segregation analyses

We tested segregation of rare, high risk loss of Function (LoF) and missense variants with CADD score >20 in *ABCA7*, *BDNF*, *NPY* and *NGFR* genes with clinical AD status. We tested segregation in 214 Caribbean Hispanic and 197 non-Hispanic White families with two or more affected members. Variants were defined as completely segregating if present in all affected family members and unaffected carriers must be at least five years younger than the average age of AD onset in families.

### Human brain DNA methylation measurement

The genome-wide DNA methylation profile was measured by the Infinium MethylationEPIC Kit (Illumina) on New York Brain Bank, NIA AD-FBS/NCRAD and EFIGA cohorts^62^. On the sample level quality control (QC), we have checked the control probes, sex mismatch, contamination, and genotype outlier calling to identify and remove those samples failed any of these QC metrics. On the CpG probe level QC, we kept those CpG sites with detection *P* value < 0.01 across all the qualified samples and mask those sample specific CpG site with new detection *P* > 0.01^111^. We further removed those CpG sites reported to have cross-hybridization problems^112,113^ and those polymorphic CpG sites^113,114^. We further corrected the dye bias for all the qualified CpG probes. For this study, we included 20 Hispanic samples with both DNA methylation and GWAS data.

### Single nucleotide polymorphism (SNP) measurement, mQTL analyses, Braak stage associations

We measured the genotype of SNP across the genome using the Infinium Global Screening Array-24 v3.0 Kit (Illumina) with the same extracted DNA samples used for the DNA methylation measurement. We have conducted the SNP level QC by removing those with missing value >5% and failed Hardy-Weinberg Equilibrium (HWE) test. As a result, there remained 4 SNPs at *ABCA7* loci, which were included into the mQTL analysis. For each of the 4 extracted SNPs at *ABCA7* loci, we conducted generalized linear regression model by treating the methylation of CpG sites at *ABCA7*, *NPY*, *BDNF*, and *NGFR* as dependent variables and genotypes of the 4 SNPs as independent variables with the adjustment of age, sex, ethnicity, and technical covariates. Those CpG sites with P value < 0.05 were considered as significant loci. For association between brain RNA expression and Braak stage, we conducted generalized linear regression model BRAAK∼Age+Gender+Gene_A+Gene_B+ Gene_A*Gene_B+ RACE+cohorts+PMI+RIN) with binary Braak stage categories (Braak 0-4 and Braak 5-6) with the adjustment of age, sex, ethnicity, post mortem interval (PMI) and RIN number for RNA quality as technical covariates. New York Brain Bank, NIA-LOAD/NCRAD and Mayo Clinic Jacksonville cohorts^62^ were used.

### Human brain sections and immunohistochemistry

Human brain sections from BA9 prefrontal cortex were also obtained from the New York Brain Bank at Columbia University (two individuals with Braak stage IV/V, two individuals with Braak stage I/II) and immunohistochemical staining for NPY was performed as described^15,115^.

## Author contributions

Conceived and designed the study: C.K. Generated the knockout line and performed the zebrafish experiments and data analyses: H.T., E.Y., P.B., N.N., N.K., R.M.D., C.K. Single cell transcriptomics: E.Y., P.B., M.I.C., C.K. Postmortem human brains: D.F., A.F.T. Performed human transcriptome and genome data analyses: Y.M., X.W., O.I., T.I.G., B.N.V., Z.Y., G.T., N.E-T., R.M. Analyzed and interpreted the human and zebrafish data: P.B., E.Y., Y.M., X.W., O.I., T.I.G., D.R-D, Z.Y., G.T., B.N.V., N.E-T., R.M., C.K. Funding: N.E-T,. R.M., C.K. Wrote the manuscript: C.K. Edited the manuscript: E.Y., P.B., X.W., O.I., B.N.V., G.T., N.E-T., R.M., C.K. All authors approved the final version of the manuscript.

## Acknowledgements

We would like to thank Taub Institute for Research on Alzheimer’s Disease and the Aging Brain Imaging Platform (USA), and Molecular Pathology platform of the Columbia University Herbert Irving Comprehensive Cancer Center (USA) for help, and New York Brain Bank (USA) for post-mortem human brain sections. We thank the contributors, who collected samples used in this study. We thank the patients and families for their participation, without whom these studies would not have been possible. This work was supported by Columbia University Schaefer Research Scholar Award, Thompson Family Foundation Program for Accelerated Medicines Exploration in Alzheimer’s Disease and Related Disorders of The Nervous System (TAME-AD), and Taub Institute Grants for Emerging Research (TIGER) (C.K.); by National Institute on Aging R01 AG067501 (Genetic Epidemiology and Multi-Omics Analyses in Familial and Sporadic Alzheimer’s Disease Among Secular Caribbean Hispanics and Religious Order) (to R.M, G.T., B.N.V as PIs and C.K. as co-investigator), and the National Institute on Aging Alzheimer’s disease Family Based Study (U24AG056270) to R.M, and B.N.V and C.K. as co-investigators. This work was also supported by National Institute on Aging [U01 AG046139, U19 AG074879, and R01 AG061796 to N.E.-T.] and Alzheimer’s Association Zenith Awards (N.E.-T.). The National Institute on Aging-AD Family-based study (NIA AD-FBS; https://www.neurology.columbia.edu/research/research-centers-and-programs/national-institute-aging-alzheimers-disease-family-based-study-nia-ad-fbs) collected the samples used in this study and is supported by National Institute on Aging (NIA) grants U24AG026395, U24AG021886, R01AG041797, and U24AG056270. Additional families were contributed to the NIA-AD FBS through NIH grants: R01AG028786, R01AG027944, RO1AG027944, RF1AG054074, U01AG052410. The NIA-AD FBS began in 2003 with the goal of recruiting large, multiply-affected families with late-onset Alzheimer’s disease (AD) for genetic research. The study created a resource of well-characterized families with late-onset AD. The initial phases of the Alzheimer’s Disease Sequencing Project (ADSP) included genotyping of hundreds of participants from NIA-AD FBS. The ADSP Follow-Up Study heavily engages resources provided by the NIA-AD FBS and depends upon the longitudinal follow-up of families, and the collection of additional families, in particular those from diverse populations. Samples include biological materials for genome wide association studies (GWAS) and whole genome sequencing (WGS), peripheral blood mononuclear cells (PBMC) for stem cell modeling, plasma for studies of metabolomics, proteomics, and biomarker research, and brain autopsy materials for bulk RNA sequencing.

Estudio Familiar de Influencia Genetica en Alzheimer (EFIGA) is a study of sporadic and familial Alzheimer’s Disease among Caribbean Hispanics recruited from clinics in the Dominican Republic and New York (R01 AG067501). The goal of this study is to identify genetic variants that increase late onset Alzheimer disease risk in this ethnic group. This study was initiated in 1998 and recruited individuals and their families in New York as well as from clinics in the Dominican Republic. Recruitment for the EFIGA began in 1998, to study the genetic architecture of AD in the Caribbean Hispanic population. Patients with familial AD were recruited and if a sibling of the proband had dementia, all other living siblings and available relatives underwent evaluation. Cases were defined as any individual meeting NINCDS-ADRDA criteria for probable or possible AD.

The results published here are in whole or in part based on data obtained from the AD Knowledge Portal (https://adknowledgeportal.org). The Mayo RNAseq study data was led by Dr. Nilüfer Ertekin-Taner, Mayo Clinic, Jacksonville, FL as part of the multi-PI U01 AG046139 (MPIs Golde, Ertekin-Taner, Younkin, Price) using samples from The Mayo Clinic Brain Bank. Data collection was supported through funding by NIA grants P50 AG016574, R01 AG032990, U01 AG046139, R01 AG018023, U01 AG006576, U01 AG006786, R01 AG025711, R01 AG017216, R01 AG003949, CurePSP Foundation, and support from Mayo Foundation. Study data included samples collected through the Sun Health Research Institute Brain and Body Donation Program of Sun City, Arizona, USA. The Brain and Body Donation Program is supported by the NINDS (U24 NS072026, National Brain and Tissue Resource for Parkinson’s Disease and Related Disorders); the NIA (P30 AG19610, Arizona Alzheimer’s Disease Core Center); the Arizona Department of Health Services (contract 211002, Arizona Alzheimer’s Research Center); the Arizona Biomedical Research Commission (contracts 4001, 0011, 05-901, and 1001, to the Arizona Parkinson’s Disease Consortium); and the Michael J. Fox Foundation for Parkinson’s Research. Study data were provided by the Rush Alzheimer’s Disease Center, Rush University Medical Center, Chicago. Data collection was supported through funding by NIA grants P30AG10161 (ROS), R01AG15819 (ROSMAP; genomics and RNAseq), R01AG17917 (MAP), R01AG30146, R01AG36042 (5hC methylation, ATACseq), RC2AG036547 (H3K9Ac), R01AG36836 (RNAseq), R01AG48015 (monocyte RNAseq) RF1AG57473 (single nucleus RNAseq), U01AG32984 (genomic and whole exome sequencing), U01AG46152 (ROSMAP AMP-AD, targeted proteomics), U01AG46161(TMT proteomics), U01AG61356 (whole genome sequencing, targeted proteomics, ROSMAP AMP-AD), the Illinois Department of Public Health (ROSMAP), and the Translational Genomics Research Institute (genomic). Additional phenotypic data can be requested at www.radc.rush.edu. The results published here are in whole or in part based on data obtained from the AD Knowledge Portal (https://adknowledgeportal.org/). These data were generated from postmortem brain tissue collected through the Mount Sinai VA Medical Center Brain Bank and were provided by Dr. Eric Schadt from Mount Sinai School of Medicine.

## Supplementary Information

Supplementary Figures 1-9

Supplementary Data 1-8

Supplementary Tables 1-5

**Supplementary Data 1**: Raw data for quantification graphs. Related to Figure 1 and Figure 4.

**Supplementary Data 2:** Gene ontology terms for biological processes in every cell cluster when *abca7* knockout animals are compared to wild type animals. Related to Figure 2.

**Supplementary Data 3:** Differentially expressed genes in *abca7* versus wild type with amyloid toxicity. Related to Figure 2 and Figure 3.

**Supplementary Data 4**: List of genes expressed in every cell cluster. Related to Figure 2 and 3.

**Supplementary Data 5:** Analyses on Mayo Clinic AD cohorts for weighed co-expression, genetic association, differential gene expression, and GO term modules. Related to Figure 4.

**Supplementary Data 6:** NicheNet analyses on AD cohorts with NPY and its receptors. Related to Figure 4.

**Supplementary Data 7:** mQTL summary for epigenetic regulatory variants for *ABCA7, NPY, NGFR*, and *BDNF*. Related to Figure 4.

**Supplementary Data 8:** Cell sorting gating strategies and raw report for all samples.

**Supplementary Figure 1:**
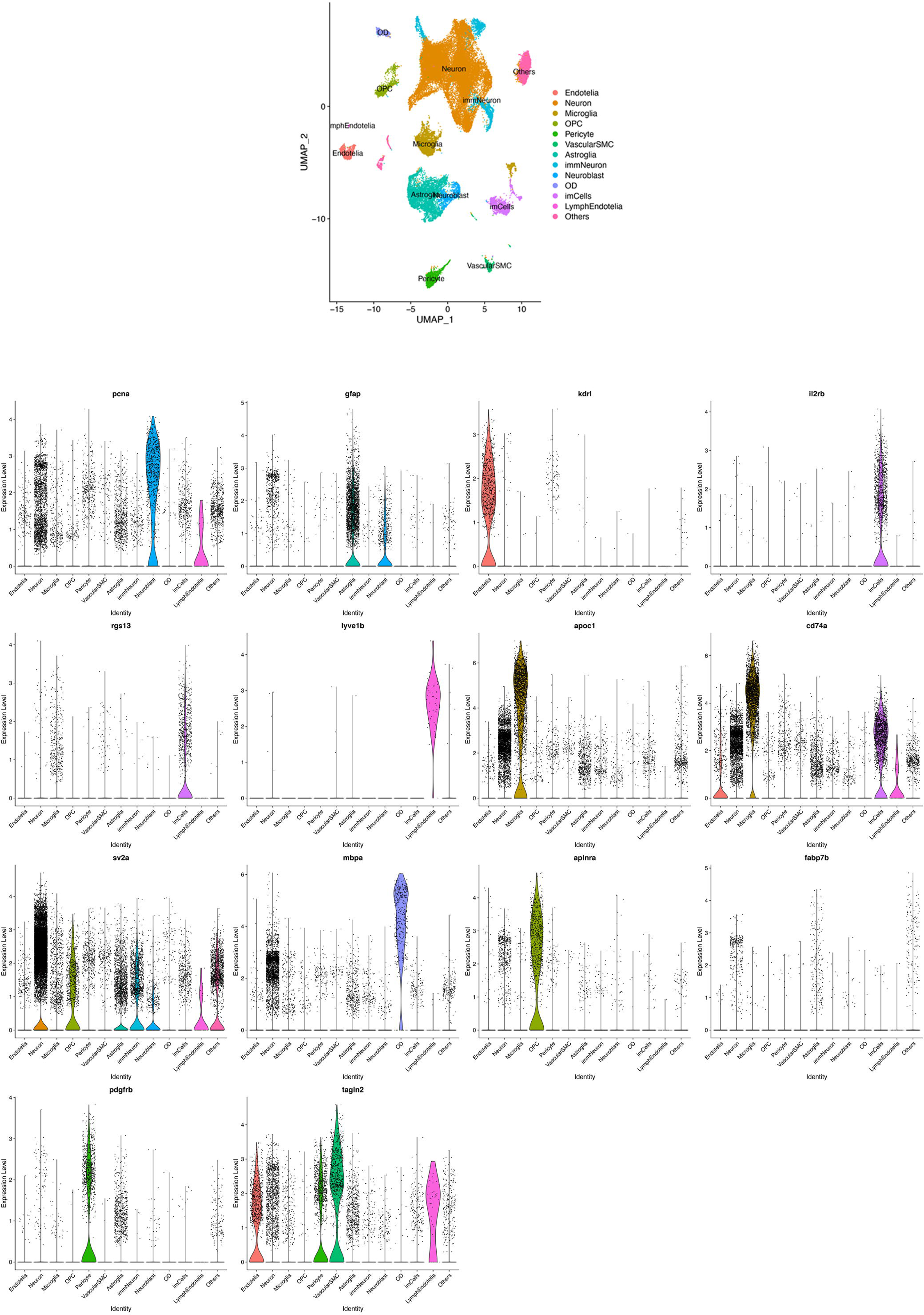
**Expression of cell specific markers for cell type identification in single cell transcriptomics**. UMAP cluster plot with cell types and violin plots for cell type marker gene expression for *pcna* (proliferating cells), *gfap* (astroglia), *kdrl* (endothelia), *il2rb* and *rgs13* (immune cells), *lyve1b* (lymphatic endothelia), *apoc1* and *cd74a* (microglia), *sv2a* (neurons), *mbpa* (oligodendrocytes), *aplnra* (oligodendrocyte progenitor cells), *fabp7b* (neuroepithelial lineage), *pdgfrb* (pericytes), *tagln2* (vascular smooth muscle cells) are shown.

**Supplementary Figure 2:**
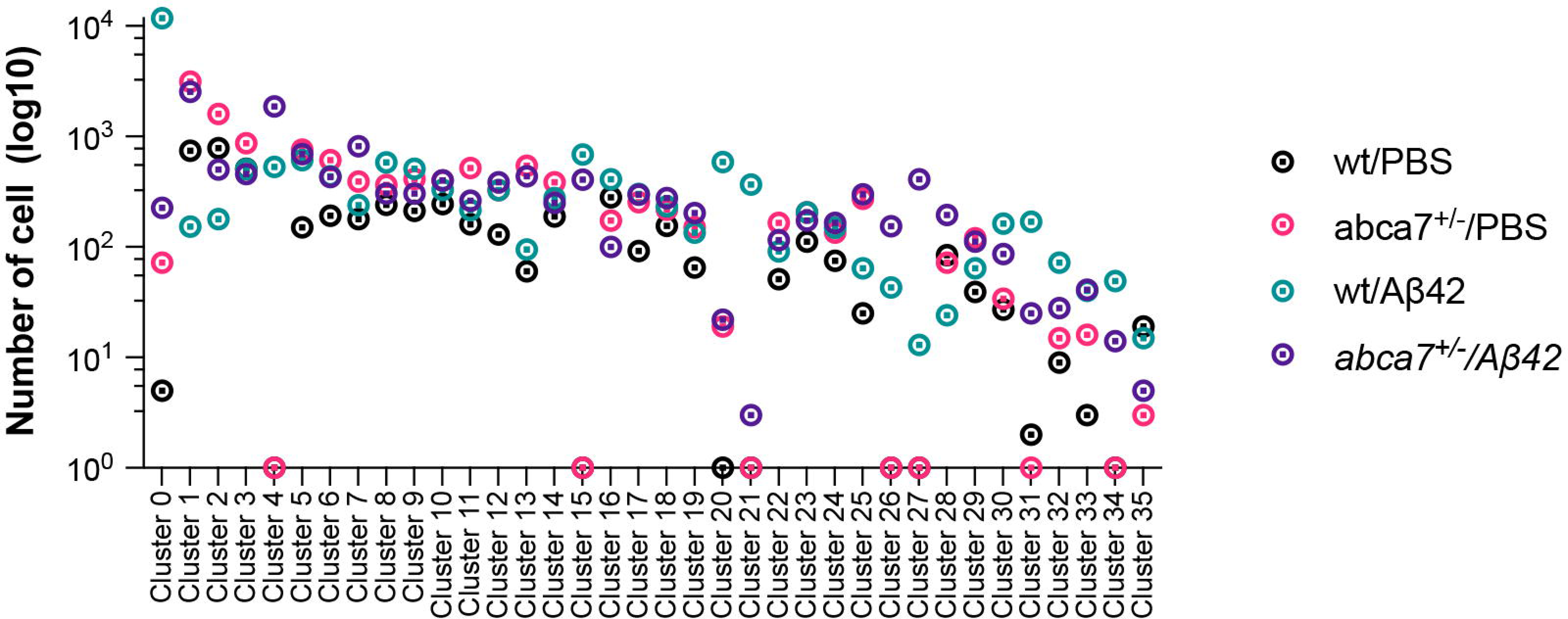
Number of cells in single cell transcriptpmics in zebrafish. The number of cells per cluster in all four experimental sets: wild type and *abca7* knockout with or without amyloid beta 42 are shown.

**Supplementary Figure 3:**
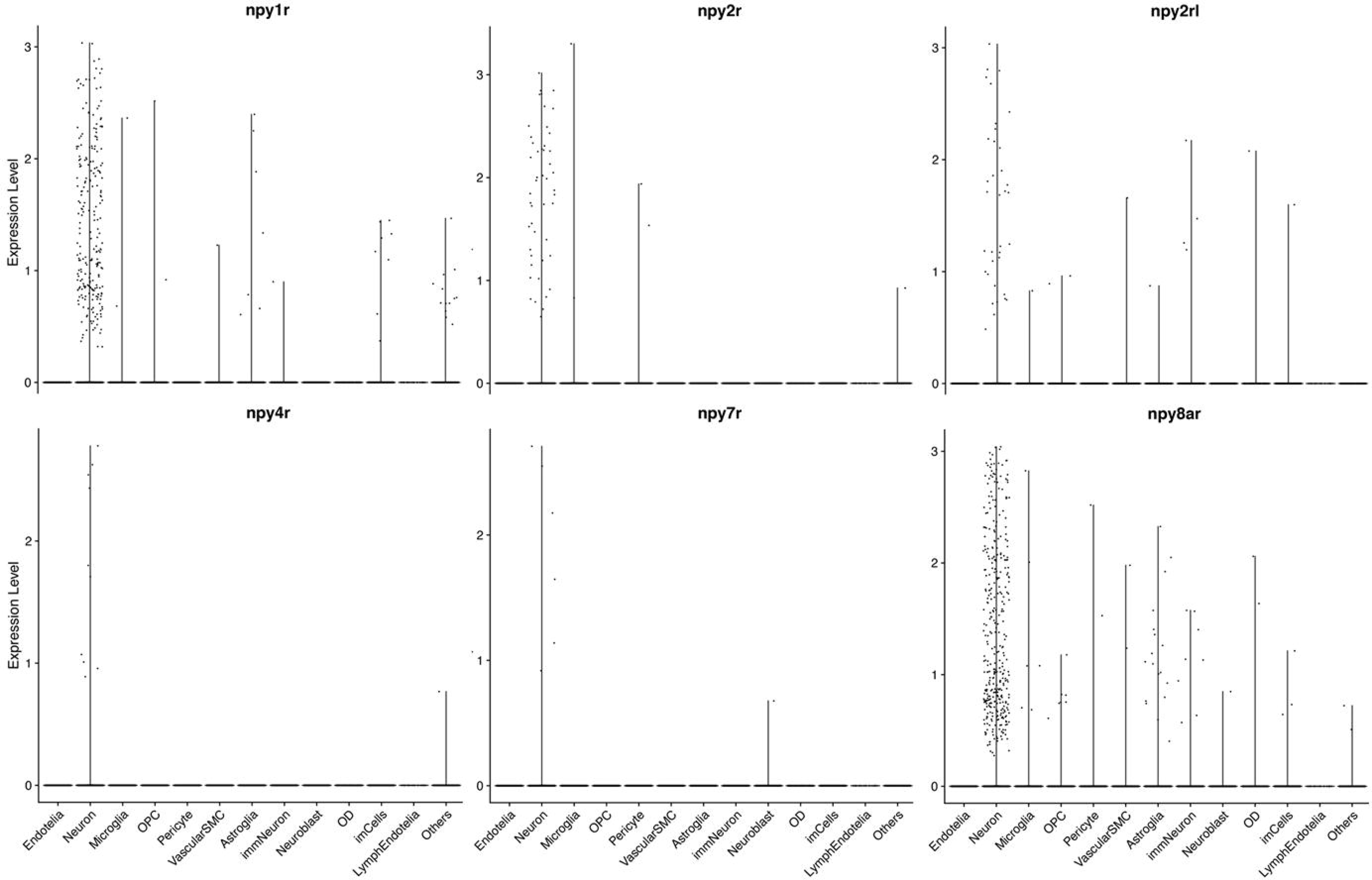
Expression plots for Npy receptors in zebrafish. Violin plots for *npy1r, npy2r, mpy2rl, npy4r, npy7r,* and *npy8ar* are shown.

**Supplementary Figure 4:**
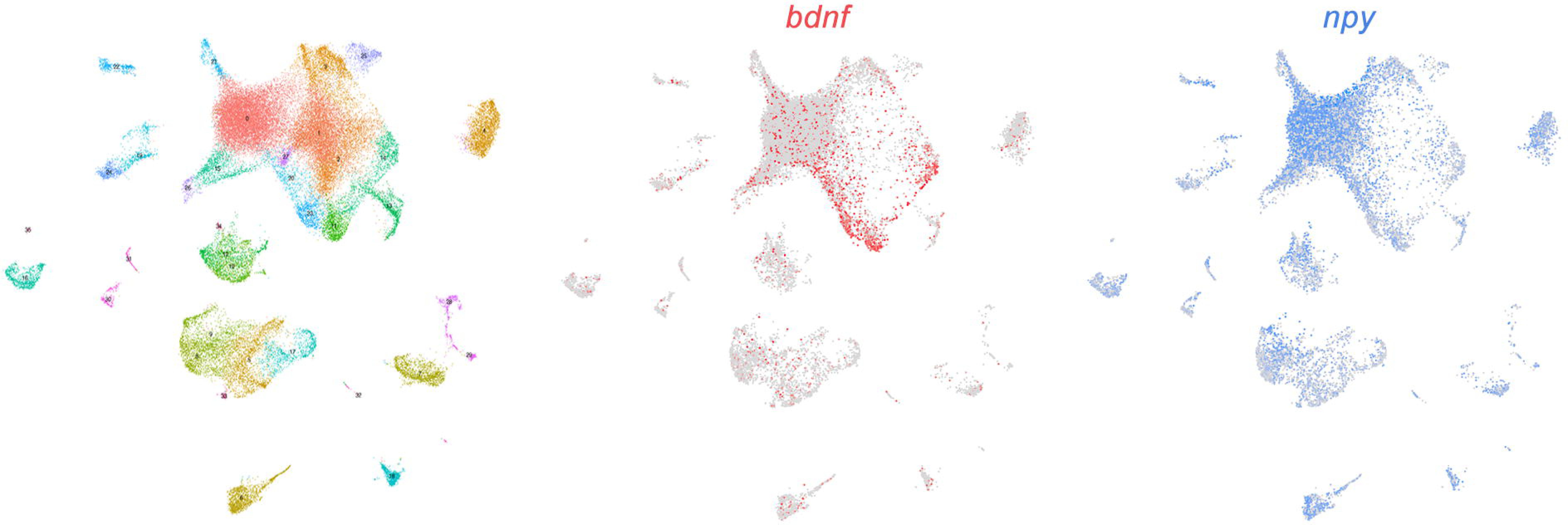
**Co-expression of *npy* and *bdnf* in zebrafish**. Total UMAP clusters and feature plots for *bdnf* (red) and *npy* (blue) expression are shown.

**Supplementary Figure 5:**
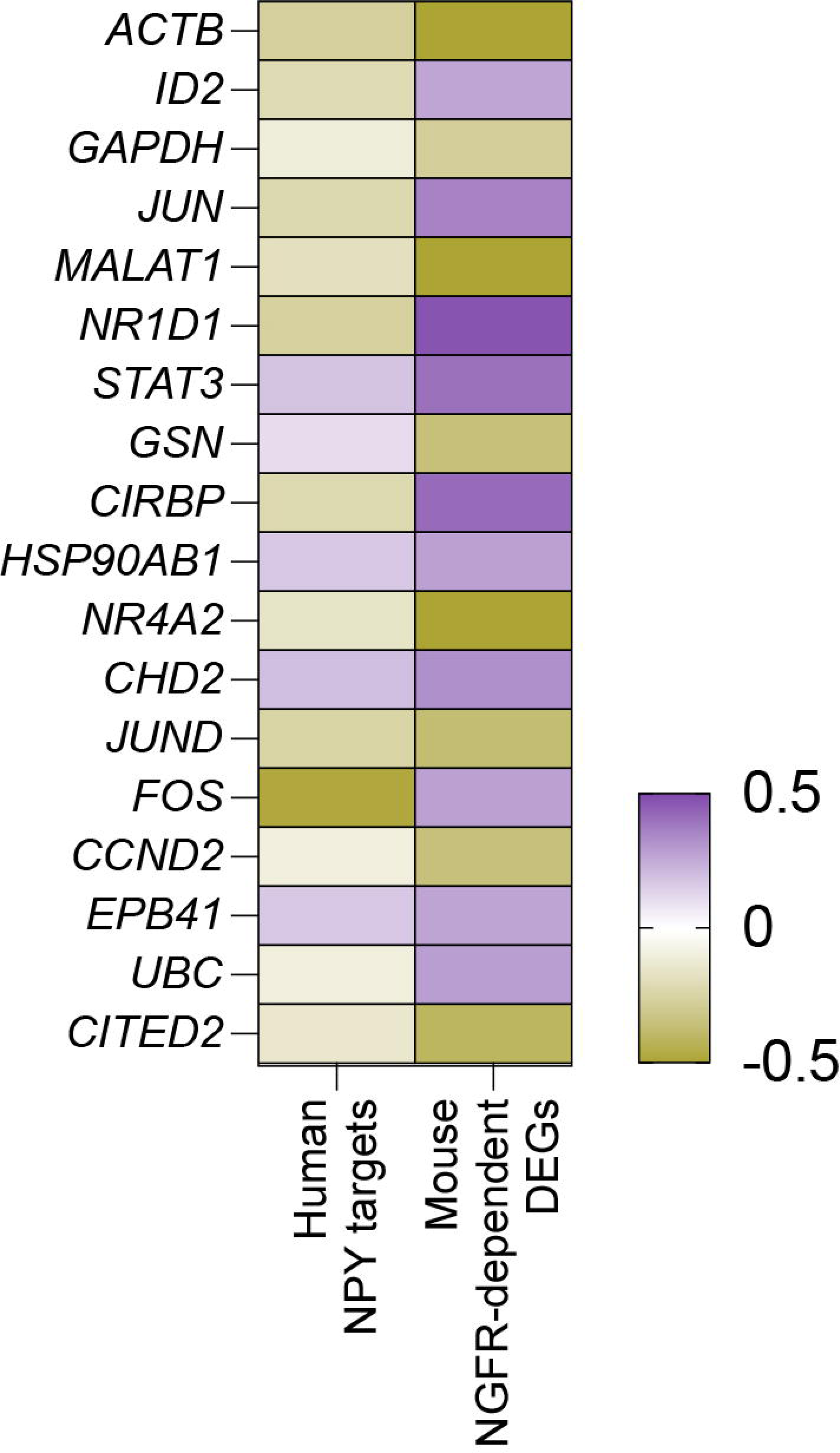
Potential common targets of NPY and NGFR in human and mouse. Comparison of human targets of NPY and differentially expressed genes after NGFR expression in mouse astroglia are shown. Potential NPY targets in humans detected with NicheNet analyses overlaps with differentially expressed genes after ectopic expression of NGFR in APP/PS1dE9 mouse model of AD.

**Supplementary Figure 6:**
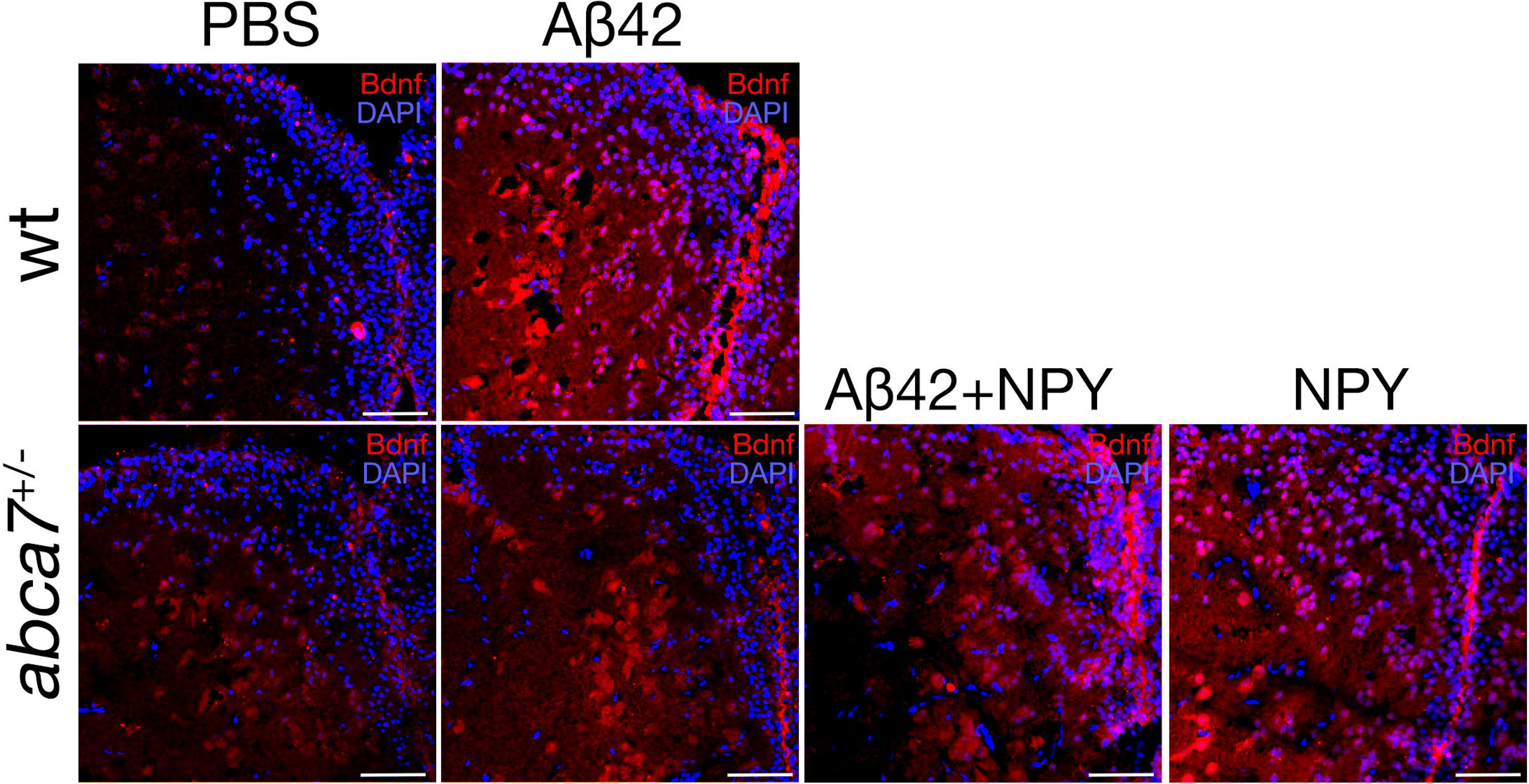
NPY can rescue the impaired BDNF expression in *abca7* knockout zebrafish after amyloid toxicity. Bdnf immunoreactivity in wild type and *abca7* knockout zebrafish with and without amyloid toxicity. Amyloid injection induced BDNF (red) in wild type animals but not in *abca7* knockout. Injection of NPY with or without amyloid beta 42 restores the induced BDNF expression levels.

**Supplementary Figure 7:**
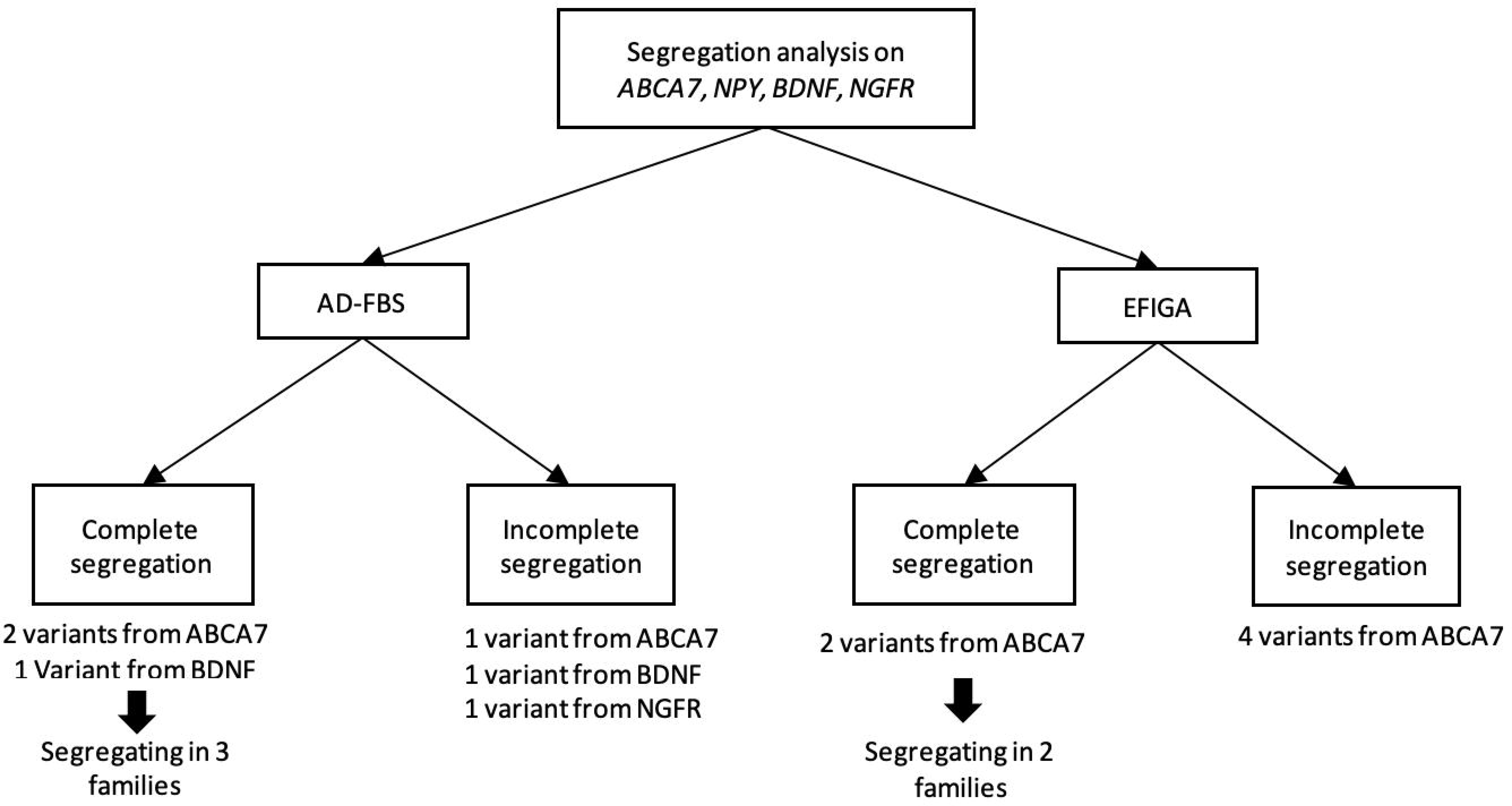
**Flowchart depicting the segregation of *ABCA7*, *NPY*, *BDNF* and *NGFR* in the Hispanic (EFIGA) and the white, non-Hispanic (AD-FBS) ancestries.**

**Supplementary Figure 8:**
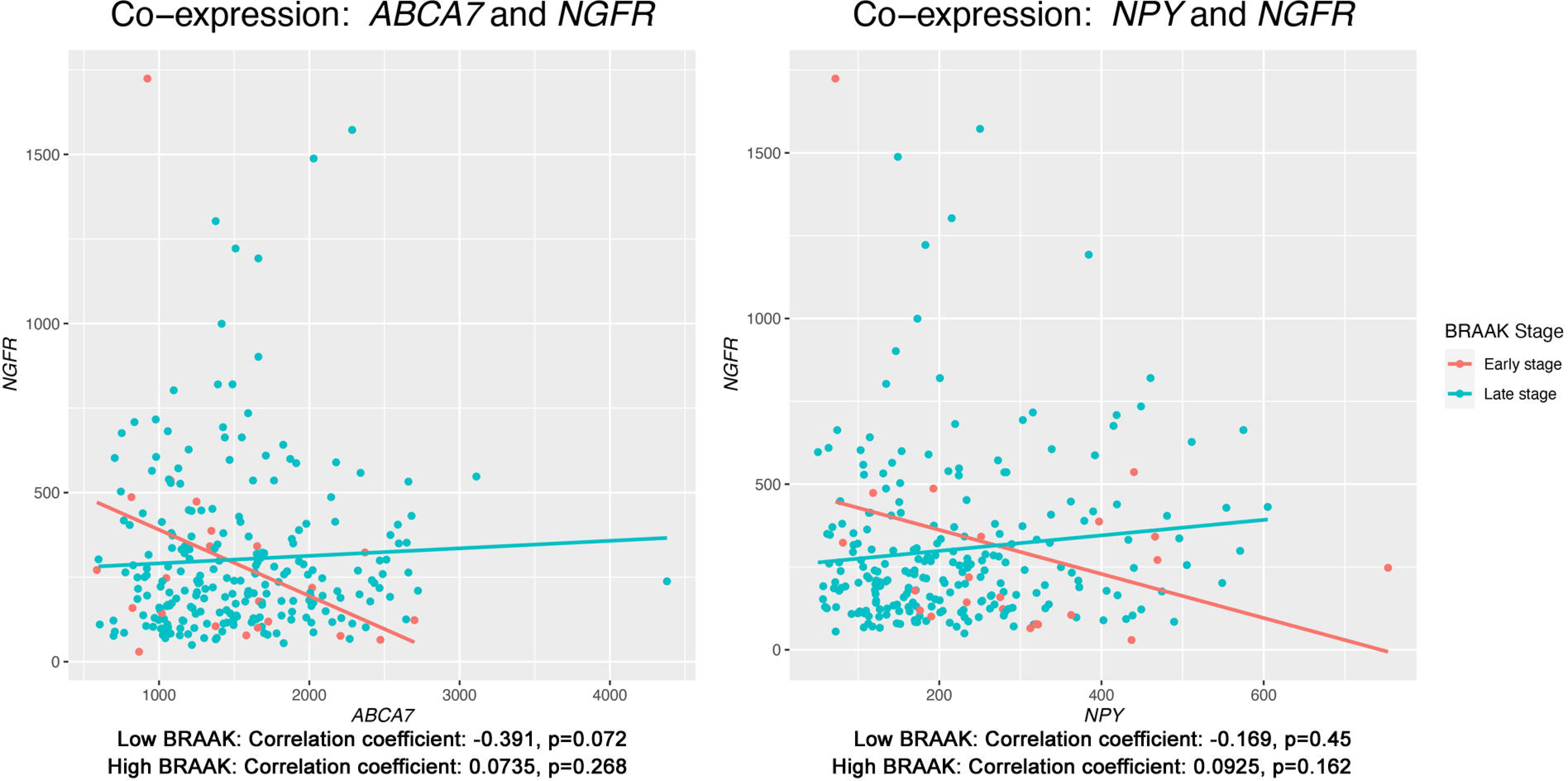
Co-expression of *ABCA7, NPY* and *NGFR* in humans in relation to the Braak stage. Plots indicate the association of expression levels of *ABCA7* and *NPY* in relation to *NGFR* in low (Braak 0-4, early stage) and high (Braak 5-6, late stage) AD. New York Brain Bank and Mayo Clinic Jacksonville cohorts were used for the analysis.

**Supplementary Figure 9:**
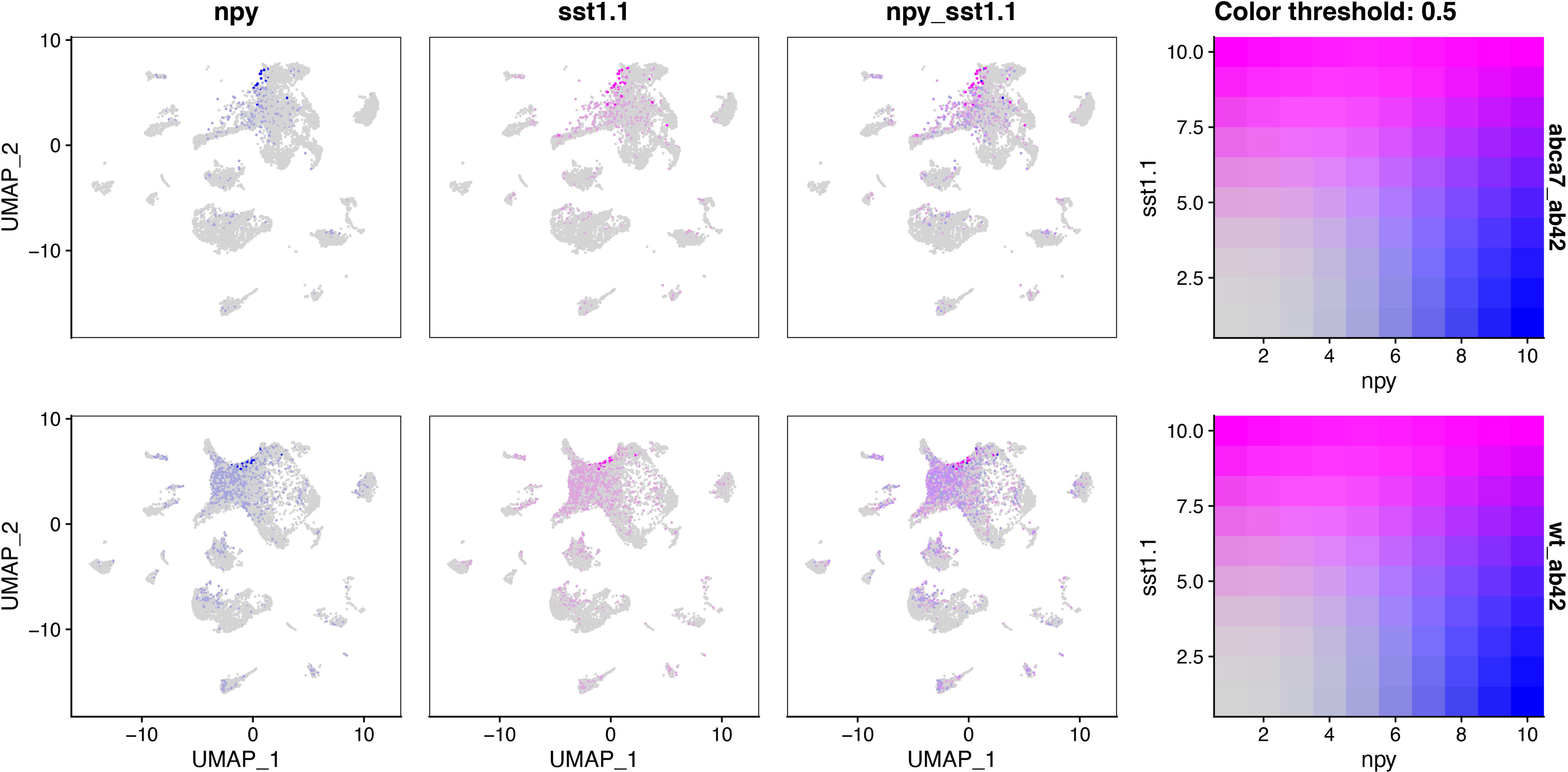
Co-expression plot for *npy* and *sst1*.*1* in zebrafish. Feature plots for *npy* (blue) and *sst1.1* (magenta) in wild type and *abca7* knockout zebrafish with amyloid injection.

**Supplementary Table 1:** Materials and reagents used in this study. Related to all figures.

**Supplementary Table 2:** Changes in the expression levels of *ABCA7, NPY* and *NPY1R* in AMP AD Mayo-TCX, Mayo-CER and ROSMAP cohorts. Related to Figure 4.

**Supplementary Table 3:** Demographic characteristics of AD-FBS and EFIGA cohorts. Related to Figure 4.

**Supplementary Table 4:** Completely segregating variants in the AD families. Related to Figure 4.

**Supplementary Table 5:** Logistic regression model on binary association of *ABCA7, BDNF, NPY,* and *NGFR* expression with Braak stages in NIA-LOAD and EFIGA cohorts. Related to Figure 4.

